# A widespread proteinaceous sulfur storage compartment in bacteria

**DOI:** 10.1101/2023.06.21.545984

**Authors:** Robert Benisch, Michael P. Andreas, Tobias W. Giessen

## Abstract

Intracellular compartmentalization is essential for all cells and enables the regulation and optimization of metabolism^1^. One of the main functions of subcellular compartments is the storage of nutrients^2–4^. As bacteria do generally not possess membrane-bound organelles, they often have to rely on functionally analogous protein-based compartments^2,5–7^. Encapsulin nanocompartments are one of the most prevalent protein-based compartmentalization strategies found in prokaryotes^5,8^. Here we show that desulfurase encapsulins represent a novel sulfur storage compartment in bacteria able to sequester large amounts of crystalline elemental sulfur. We determined the 1.78 Å cryo-EM structure of a 24 nm desulfurase-loaded encapsulin highlighting the molecular details of the protein shell and desulfurase encapsulation. We found that elemental sulfur crystals can be formed inside encapsulin shells in a desulfurase-dependent manner with L-cysteine acting as the sulfur donor. Intracellular sulfur accumulation can be influenced by the concentration and type of sulfur source in growth media. The selectively permeable protein shell allows the long-term intracellular storage of redox-labile elemental sulfur by excluding cellular reducing agents from its interior. We found that encapsulation substantially improves desulfurase activity and stability while also preventing substrate inhibition. These findings represent the first example of a dedicated and widespread storage system for the essential element sulfur in bacteria and provide the basis for understanding how this novel protein-based storage compartment is integrated within bacterial metabolism.

## Introduction

Subcellular compartmentalization is essential for all cells and enables the regulation and optimization of metabolism^1,6^. This is not only true for large and complex eukaryotic cells, but also for prokaryotes. In recent years, significant progress has been made to highlight that bacterial cells are highly organized entities often relying on protein-based strategies to coordinate and compartmentalize complex metabolic functions^6,9–13^. One of these strategies are protein organelles and compartments which represent nano-sized functional analogues of eukaryotic membrane organelles and utilize semipermeable protein shells to sequester specific enzymes and processes. For example, bacterial microcompartments (BMCs) sequester combinations of enzymes in self-assembling protein shells and are involved in the anabolic fixation of carbon^14,15^ and catabolic processes like carbon and nitrogen source utilization^7,16^. Besides serving as nanoscale reaction chambers, another important use of protein compartments is the storage of nutrients^1,6,10,13^. The most widely distributed protein-based storage system is ferritin, an 8-12 nm protein cage used by eukaryotic and prokaryotic cells to store iron^2^. Many cells contain further systems for storing nutrients such as polyphosphate-^17^, polyhydroxyalkanoate-^18^, and sulfur-storage granules or globules whose detailed functions, compositions, and formation are still being debated^13,19^. In general, storage compartments enable organisms to accumulate and retain high-value compounds for later use when encountering changing, nutrient-limited, or stress conditions^17,18,20^.

A further and only recently discovered class of prokaryotic protein compartments involved in storage and other functions are encapsulin nanocompartments (encapsulins)^5,21^. Encapsulins consist of self-assembling protein shells sequestering dedicated cargo enzymes and are among the most widespread protein compartments in prokaryotes^5,8,22^. Cargo encapsulation is mediated by targeting sequences present at the N- or C-terminus of all cargo proteins^5,21,23,24^. Encapsulin shells possess icosahedral symmetry with triangulation numbers of T=1 (60 subunits, 24 nm), T=3 (180 subunits, 32 nm), or T=4 (240 subunits, 42 nm) and an evolutionary connection with viral capsids has been proposed^5,21, 25–27^. Encapsulins are classified into four families based on sequence similarity and operon organization, with Family 1 encapsulins having been shown to be involved in iron storage, detoxification, and stress resistance^8,22, 26–29^. Bioinformatic analyses have further identified a novel widespread Family 2 encapsulin system putatively involved in redox or sulfur metabolism^8^. A recent study in *Synechococcus elongatus* confirmed that this Family 2A system is induced under sulfur starvation conditions and encodes a cysteine desulfurase (CD) cargo protein sequestered inside an encapsulin shell^23^. CDs are pyridoxal-5’-phosphate (PLP)-dependent enzymes that catalyze the desulfurization of L-cysteine, yielding L -alanine and an enzyme-bound persulfide intermediate^30,31^. It was found that desulfurase cargo loading is facilitated by an N-terminal cargo-loading domain (CLD) and that desulfurase activity is increased upon encapsulation^8,23^. So far, the molecular logic of CD encapsulation as well as the biological function of this class of encapsulins are unknown.

Here, we present structural and biochemical data on a cysteine desulfurase encapsulin system found in *Acinetobacter baumannii* 118362, a member of the *Acinetobacter calcoaceticus*/*baumannii* complex. Using cryo-electron microscopy (cryo-EM), we determine the 1.78 Å structure of the Family 2A encapsulin shell and report evidence for a novel cargo-loading mechanism. We find that encapsulation increases CD stability and notably enables high catalytic activity in the absence of a sulfur acceptor. CD activity can lead to the mineralization of large amounts of crystalline elemental sulfur inside the encapsulin shell which is protected from the reducing environment of the cytosol. Together, our data suggest that desulfurase encapsulin systems represent a novel and widespread intracellular sulfur storage system in bacteria.

## Results

### Cysteine desulfurase encapsulin operons are widespread in bacteria

All encapsulin shell proteins possess the HK97 phage-like fold, a widespread viral capsid protein fold found in bacteriophages of the order Caudovirales and select eukaryotic viruses including members of the Herpesviridae^32^. Recent structural and phylogenetic analyses suggest an evolutionary relationship between encapsulins and viruses^5,8^. It has been proposed that encapsulins originate from defective prophages whose capsid protein has been co-opted by the prokaryotic cellular host to now serve its own metabolic needs and increase its fitness. Recent sequence similarity and gene neighborhood analyses allowed the classification of encapsulin systems into four families^8^. So far, mostly Family 1 encapsulins have been characterized with only one example of a Family 2 system – a CD encapsulin from *S. elongatus*^23^ – having been studied. Based on the absence or presence of a putative cNMP-binding insertion domain in the encapsulin shell protein, Family 2 encapsulins can be further classified into Family 2A and 2B, respectively^5,8^.

Phylogenetic analysis of all identified Family 2A CD encapsulins revealed that these systems are prevalent and widespread in bacteria, with 1,462 CD encapsulin operons identified across 11 bacterial phyla (Fig. 1a,b and Supplementary Data 1). Most CD encapsulins are present in Proteobacteria, Actinobacteria, Bacteroidetes, and Cyanobacteria and many of these systems can be found in important model organisms and pathogens including *Mycobacterium leprae*, *Mycobacterium avium*, *Burkholderia cepacia*, *Klebsiella pneumoniae*, and *Acinetobacter baumannii*.

**Fig. 1:**
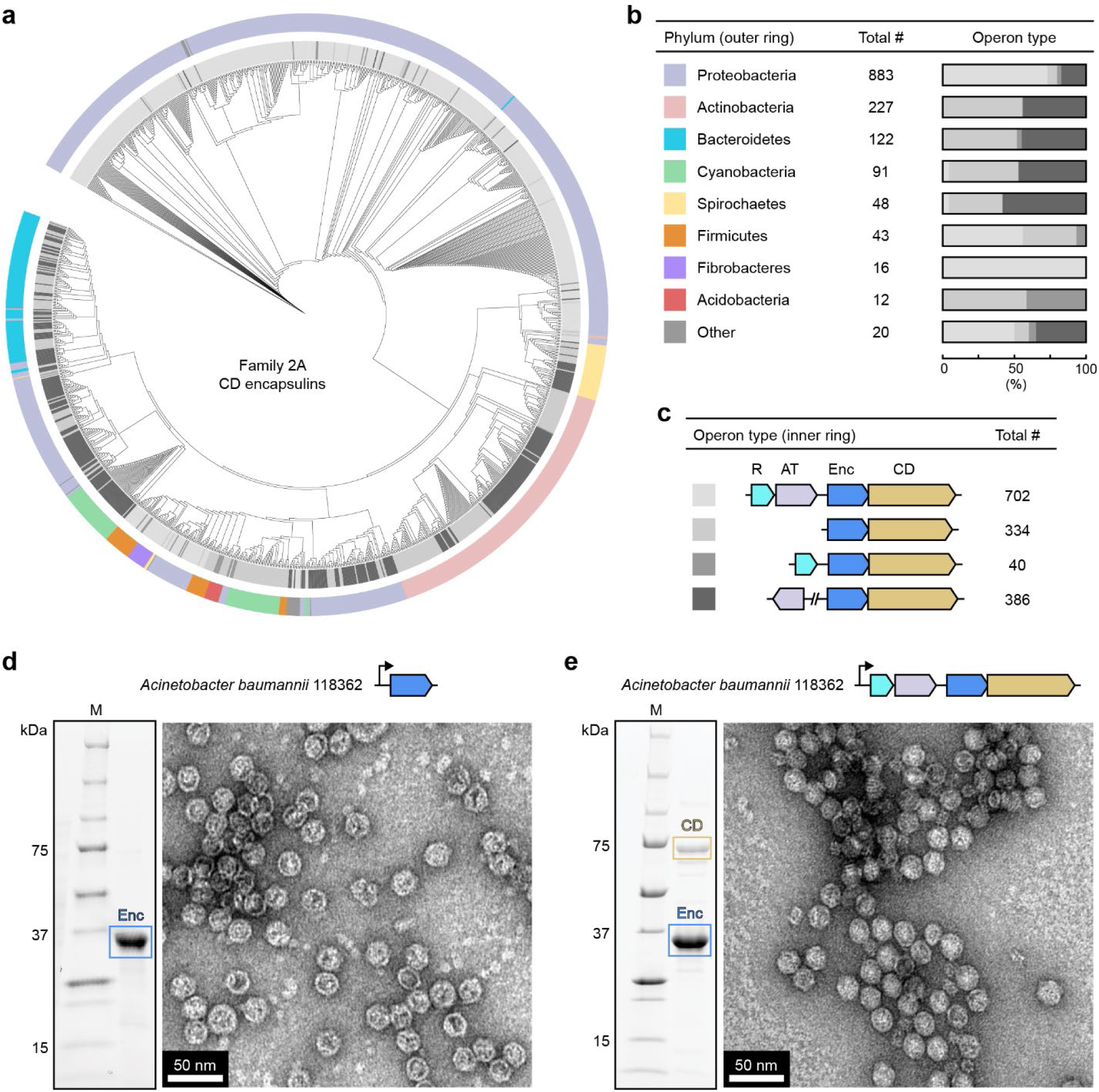
Distribution and diversity of cysteine desulfurase (CD) encapsulins. (**a**) Phylogenetic tree of 1,462 Family 2A CD encapsulins highlighting their distribution in bacterial phyla and operon type diversity. The outer ring color indicates bacterial phyla distribution (see **b**) and the gray scale inner ring highlights operon type distribution (see **c**). (**b**) Bacterial phyla encoding CD encapsulin operons and number of identified CD encapsulin operons per phylum (left and middle). Distribution of the four identified operon types (see **c**) in bacterial phyla (right). (**c**) The four identified operon organizations for CD encapsulin systems and their prevalence. R: rhodanese, AT: L-serine *O*-acetyltransferase, Enc: encapsulin shell protein, CD: cysteine desulfurase. (**d**) SDS-PAGE analysis (left) and negative stain TEM micrograph (right) of the heterologously expressed and purified *Acinetobacter baumannii* 118362 CD encapsulin shell. (**e**) SDS-PAGE analysis (left) and negative stain TEM micrograph (right) of purified CD-loaded *Acinetobacter baumannii* 118362 encapsulin resulting from the heterologous expression of a four-gene operon. M: molecular weight marker.

Closer analysis of CD encapsulin gene clusters identified four common operon organizations (Fig. 1c). All operons code for an encapsulin shell protein and a CD. Often, two additional co-regulated operon components can be present. These are annotated as a rhodanese (R) and L-serine *O*-acetyltransferase (AT). Rhodaneses are a diverse class of proteins with various functions, one of them being to serve as sulfur acceptor proteins^33–38^. Sulfur acceptors generally directly interact with CDs to facilitate the transfer of the CD-bound sulfur atom – intermittently stored as a persulfide intermediate – to a conserved cysteine residue in the acceptor protein^39^. Sulfur acceptors can then distribute sulfur to various downstream processes like iron-sulfur cluster assembly or thiocofactor biosynthesis^30,31,35^. L-serine *O*-acetyltransferases are key enzymes in the biosynthesis of L-cysteine, converting L-serine into *O*-acetyl-L-serine, the direct precursor of L-cysteine^40^. Thus, gene annotation suggests that CD encapsulin operons are involved in sulfur metabolism.

### Heterologous expression of an *Acinetobacter* desulfurase encapsulin operon identifies CD as the sole cargo protein

Here, we focus on a CD encapsulin operon found in a member of the *Acinetobacter calcoaceticus*/*baumannii* complex (*Acinetobacter baumannii* 118362) encoding all four operon components discussed above – rhodanese (J517_0525), L-serine *O*-acetyltransferase (J517_0526), encapsulin shell protein (J517_0527), and CD (J517_0528). Heterologous expression of the encapsulin shell gene or the complete native four-gene operon in *E. coli* BL21 (DE3), followed by purification via polyethylene glycol precipitation as well as size exclusion and ion exchange chromatography, yielded readily assembled encapsulin nanocompartments as confirmed by negative stain transmission electron microscopy (TEM) (Fig. 1d,e and Extended Data Fig. 1a). Consistent with previous reports, protein shells appeared spherical with a diameter of ca. 24 nm, suggesting a T=1 shell assembly^5,23^. However, only when the four-gene operon was expressed could a second major co-purifying band on SDS-PAGE be observed (Fig. 1e). Based on molecular weight and mass spectrometric analysis, this band was identified as CD (71 kDa), indicating that CD represents the sole cargo protein of this Family 2A encapsulin system. The purified sample was additionally subjected to native PAGE analysis resulting in a major 1 MDa band and no lower molecular weight bands, further confirming that CD is likely encapsulated inside the encapsulin shell (Extended Data Fig. 1b).

### Single particle cryo-EM analysis of the desulfurase-loaded encapsulin

To gain molecular level insights into the structure and cargo-loading mechanism of the CD-loaded encapsulin, single particle cryo-EM analysis was carried out (Extended Data Fig. 2). The encapsulin shell was determined to 1.78 Å via icosahedral (I) refinement (Extended Data Fig. 2a-c). This represents the highest resolution encapsulin shell structure reported to date and allowed for accurate atomic model building (Extended Data Fig. 2d,e). As suggested by negative stain TEM, the encapsulin shell was found to be 24 nm in diameter and to consist of 60 subunits, showing T=1 icosahedral symmetry (Fig. 2a). In contrast to Family 1 encapsulins, Family 2A encapsulins have been reported to possess turret-like morphology at their 5-fold vertices, similar to many HK97-fold Caudovirales capsids^5,23,32^. This is also the case for this Family 2A *Acinetobacter* encapsulin where an extended C-terminus and an extra 12° backwards tilt of the shell protein at the 5-fold symmetry axis results in turret-like vertices (Extended Data Fig. 3a,b). The asymmetric unit contains a single shell protein subunit which exhibits the canonical HK97 phage-like fold consisting of an A-domain (axial domain), P-domain (peripheral domain), and E-loop (extended loop) (Fig. 2b)^32^. Whereas Family 1 encapsulins possess an N-terminal helix located on the interior of the assembled shell^5^, this Family 2A encapsulin contains an N-arm extension reminiscent of HK97-fold viruses^23,32^. The N-arm interacts with neighboring subunits to form a chainmail-like topology (Extended Data Fig. 3c), often observed in HK97-fold viral capsids^41^, and two N-arms outline and mostly close the pore found at the 2-fold symmetry axis (Extended Data Fig. 3d). The shell contains differently sized pores at the 5-, 3-, and 2-fold axes of symmetry with likely only the 5-fold pore being large enough (6 Å) for small molecule transmission to the compartment lumen (Fig. 2c-e and Extended Data Fig. 3e). The exterior and narrowest point of the 5-fold pore are mostly uncharged and non-polar, in contrast to the Family 2A *S. elongatus* 5-fold pore, reported to be positively charged^23^. Pore size likely limits the range of molecules able to enter and exit the compartment, as has been proposed for other encapsulin systems^5,42^. The 6 Å 5-fold pore, however, is likely large enough to allow the substrate (L-cysteine) and product (L-alanine) of the encapsulated CD to pass through, whereas the 2- and 3-fold pores are likely too restrictive.

**Fig. 2:**
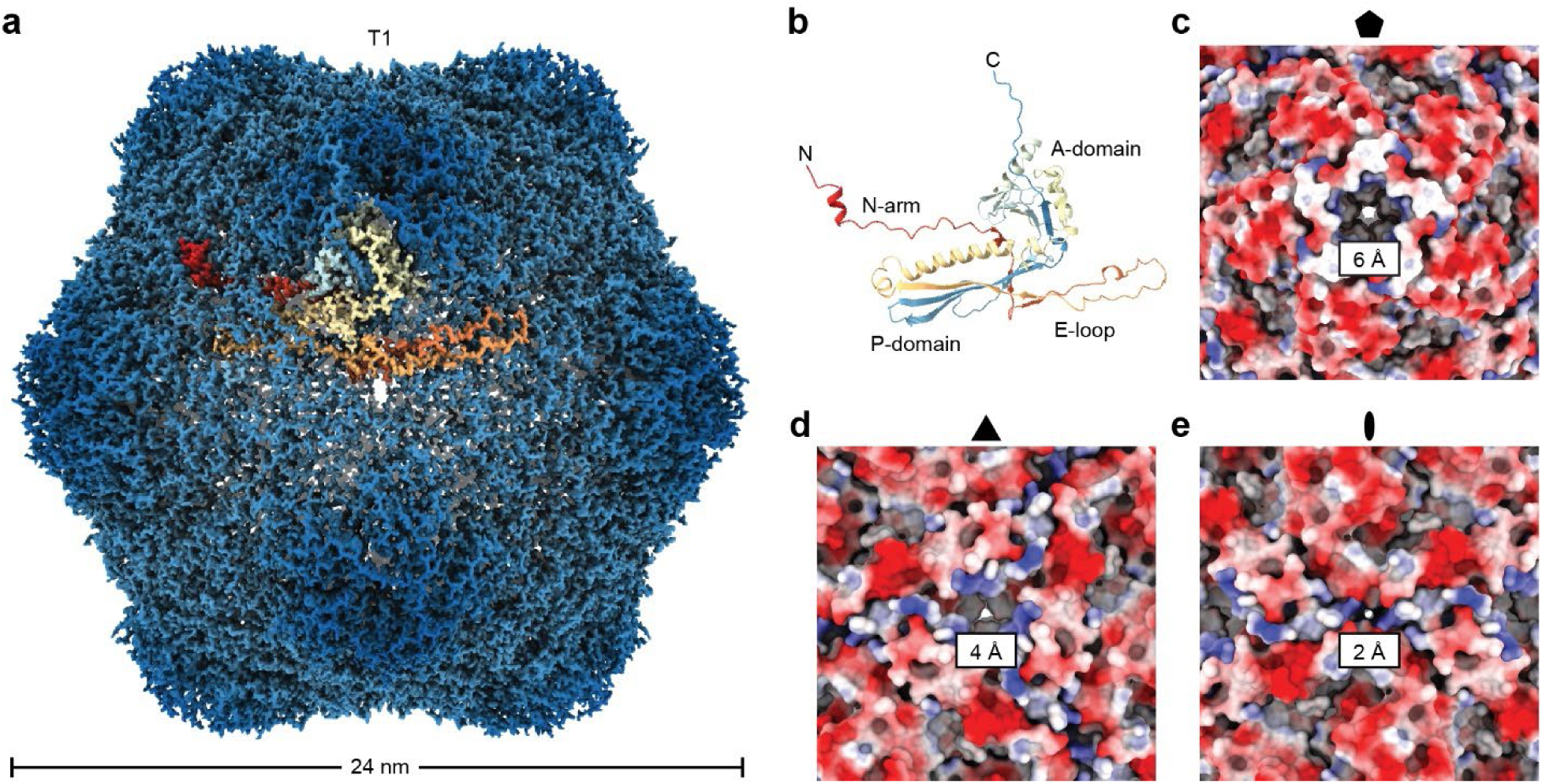
cryo-EM analysis of the encapsulin shell. (**a**) cryo-EM density resulting from icosahedral (I) symmetry refinement of the CD-loaded encapsulin. The shell is 24 nm in diameter and exhibits T=1 icosahedral symmetry. Cargo densities are not visible in the I refinement. Shell density was colored radially from the center of the shell. One subunit is highlighted in rainbow coloring from red (N-terminus) to blue (C-terminus). (**b**) A single HK97-fold encapsulin subunit is shown in rainbow coloring highlighting the canonical A (axial)-domain, P (peripheral)-domain, E (extended)-loop, and N (N-terminal)-arm. (**c**) View from the shell exterior down the 5-fold symmetry axis highlighting the 5-fold pore. Electrostatic coloring is shown. The narrowest point of the 5-fold pore is 6 Å wide. (**d**) View from the shell exterior down the 3-fold symmetry axis highlighting the 3-fold pore. Electrostatic coloring is shown. The narrowest point of the 3-fold pore is 4 Å wide. (**e**) View from the shell exterior down the 2-fold symmetry axis highlighting the 2-fold pore. Electrostatic coloring is shown. The narrowest point of the 2-fold pore is 2 Å wide.

Whereas icosahedral (I) refinement yielded the highest quality shell density, no signal for internalized CD cargo could be observed. However, C1 refinement resulted in a 2.18 Å volume where clear internal densities could be visualized (Fig. 3a). These represent the encapsulated CD cargo with 12 low-resolution densities located below each of the 12 pentameric vertices of the T=1 icosahedral shell. As has been reported for most other encapsulins^5,27,43,44^, the observed CD cargo densities are substantially lower resolution (∼15 Å) than the shell, likely due to flexible tethering to the shell interior, with only a limited number of CLD residues tightly interacting with the luminal surface (see below). The flexible linker sequences are not visible in the cryo-EM reconstruction resulting in the cargo densities appearing disconnected from the shell. CD belongs to the class II SufS/CsdA-like desulfurases with almost all characterized members of this class forming stable homodimers^31,45^. This is consistent with the observed size of the internal cargo densities and SEC analysis of the unencapsulated *Acinetobacter* CD (Extended Data Fig. 4). Thus, the maximal number of CD cargo proteins per encapsulin shell is likely 24, or 12 CD dimers which is in good agreement with cargo-loading estimates based on SDS-PAGE gel densitometry analysis (Fig. 1e). Shell density subtraction followed by 2D classification of shell-subtracted particles further confirmed the presence of cargo, yielding 2D classes with clearly visible internal densities representing CD (Fig. 3b).

**Fig. 3:**
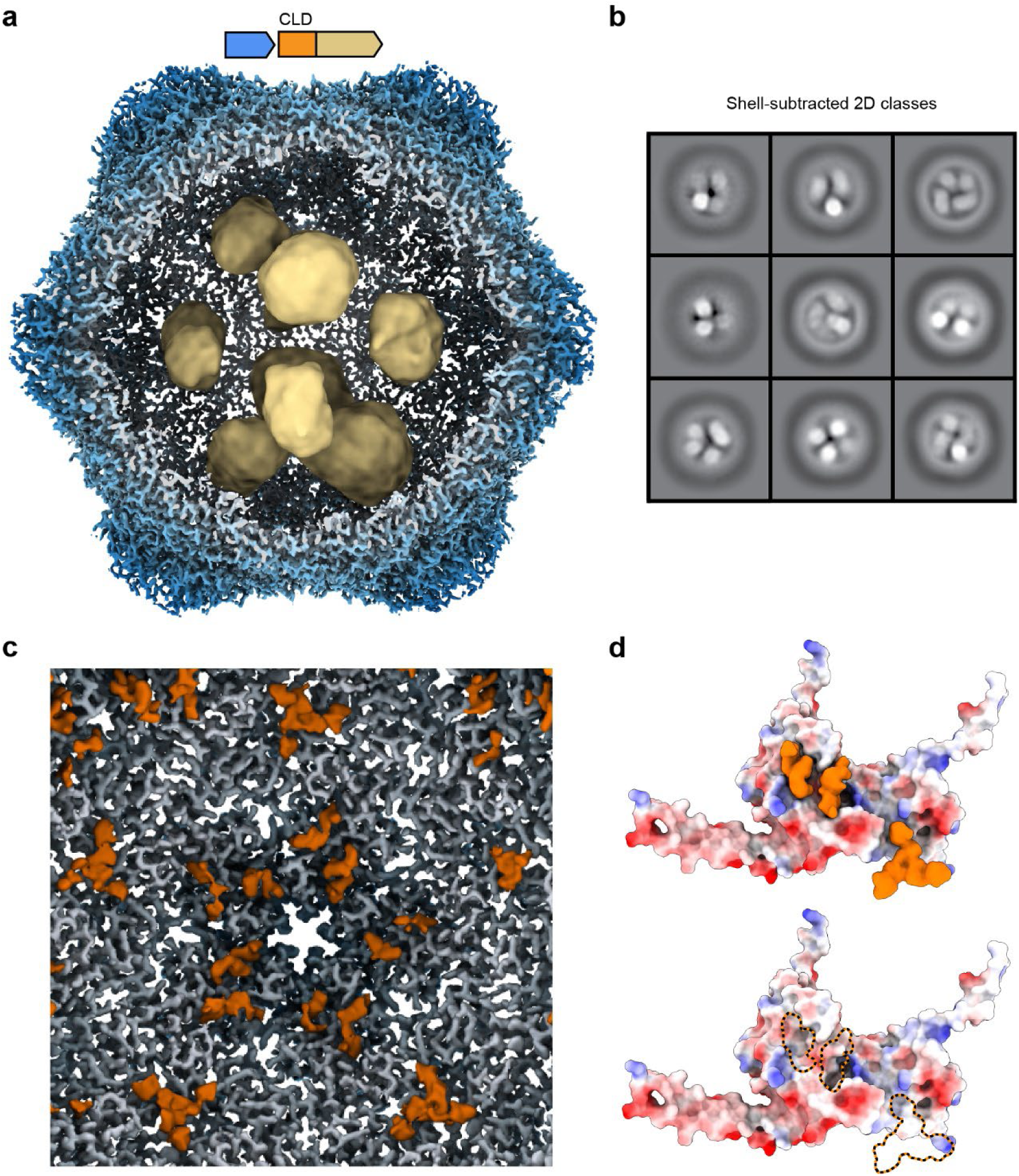
cryo-EM analysis of the CD cargo inside the encapsulin shell. (**a**) Interior view of the cryo-EM density resulting from an asymmetric (C1) refinement of the CD-loaded encapsulin showing internal CD cargo densities (yellow). The shown volume is a composite of the 2.18 Å shell density and a gaussian blurred (sdev 2, ChimeraX^46,47^) map to highlight internal lower resolution CD cargo densities. Shell density was colored radially from the center of the shell. (**b**) Shell-subtracted 2D classes of CD-loaded encapsulin highlighting discrete internal CD densities. (**c**) Composite volume of the C1 high resolution shell and a gaussian blurred map to highlight extra shell-associated densities (orange) around the 3- and 5-fold symmetry axes likely belonging to the N-terminal cargo-loading domain of the CD cargo. (**d**) A single encapsulin shell protein subunit (electrostatic coloring) is shown with (top) or without (bottom) the closely associated non-shell density (orange) likely representing the CD CLD to highlight their interaction at the shell protein A-domain (5-fold) and P-domain (3-fold). The interaction surfaces are primarily uncharged and non-polar (bottom, outlined in orange-black).

One unusual feature of CD is the presence of a ca. 225 residue long unannotated and disordered N-terminal domain, in addition to the catalytic C-terminal desulfurase domain. This domain is rich in proline, glycine, and serine and is not well conserved among putative Family 2A CD cargo proteins with only 5 relatively short motifs found to be partially conserved (Extended Data Fig. 5a)^8,23^. Previously, it was shown that a similar domain in a Family 2A system from *S. elongatus* acts as the CLD responsible for mediating cargo loading into the encapsulin shell and potentially interacts with the interior shell surface close to the 3-fold pores^23^. In our cryo-EM analysis, we observed additional non-shell densities along the interior surface of the encapsulin lumen, localized around the 3-fold pores and A-domains (Fig. 3c). These densities likely represent parts of the N-terminal CLD, however, resolution is too low for model building or sequence assignment. The observed CLD densities are not connected and localized at mostly non-polar or hydrophobic surface patches (Fig. 3d and Extended Data Fig. 5b). This suggests a model of CLD-shell interaction where different parts of the N-terminal domain specifically interact with conserved parts of the shell interior in a discontiguous way, connected by flexible linker sequences, not visible in our cryo-EM density. The 5 conserved motifs found in the CLD (Extended Data Fig. 5a) may represent the residues interacting with the shell while the long stretches of less conserved sequence between them could serve as flexible linkers. The identified surface patches at the 3-fold pores and A-domains are mostly conserved (Extended Data Fig. 5b), suggesting that the observed CLD-shell interaction may be conserved across Family 2A desulfurase encapsulins.

### CD-loaded encapsulins actively accumulate and store elemental sulfur inside cells

Closer inspection of raw cryo-EM micrographs revealed that ∼15% of shells contained large electron-dense puncta exclusively localized to the interior of encapsulin shells (Fig. 4a). Interestingly, in the absence of CD, no puncta could be observed, indicating that CD might play a role in puncta formation. High-signal puncta could also be observed inside the protein shell in a number of 2D class averages of the CD-loaded encapsulin (Fig. 4b). Subtraction of the shell signal followed by 2D classification yielded a number of 2D classes where both puncta and CD cargo densities seem to co-localize to the encapsulin lumen. The puncta are roughly 10-15 nm in diameter, occupying ∼50% of the luminal volume (Extended Data Fig. 6a).

**Fig. 4:**
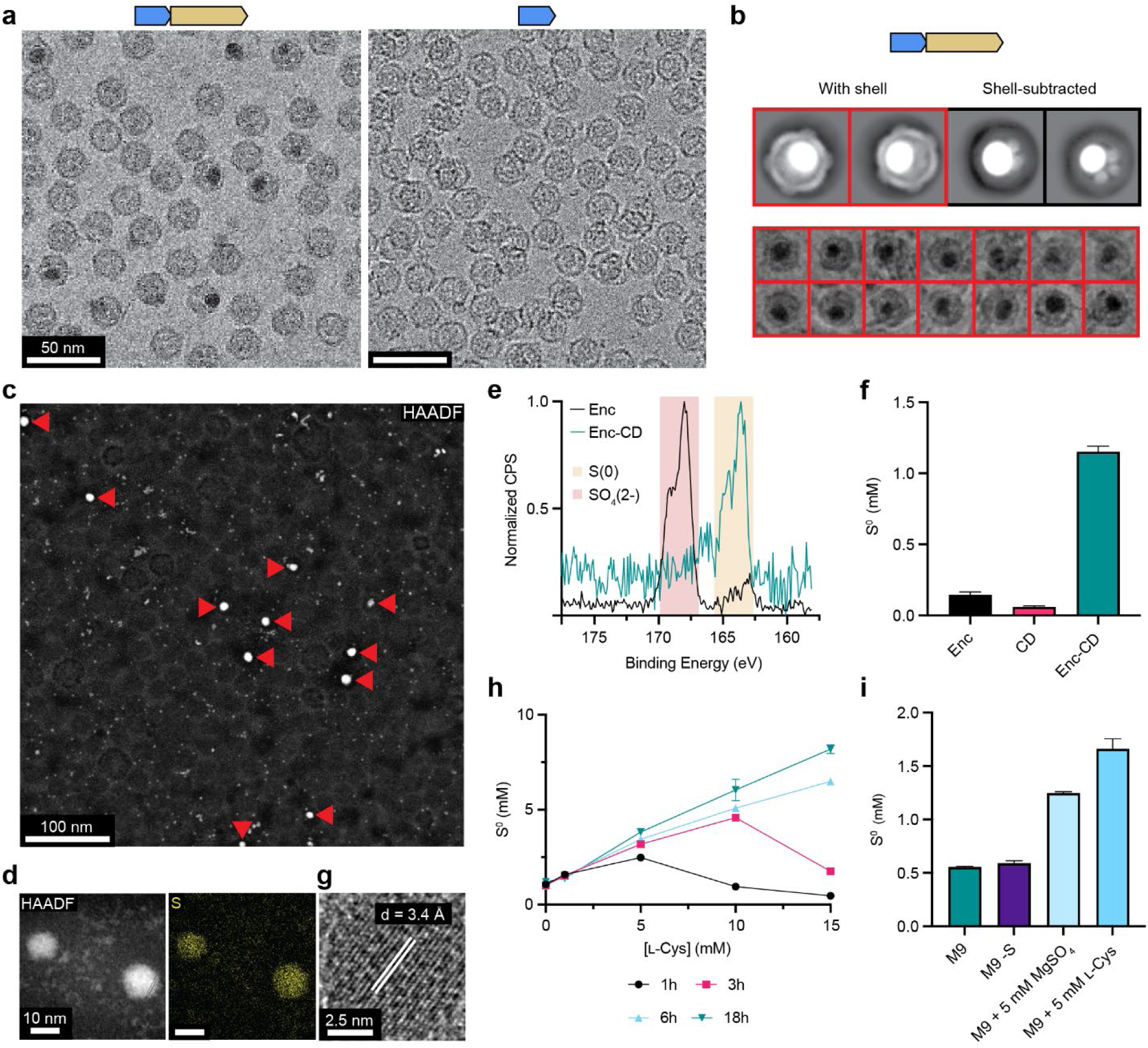
Identification and characterization of elemental sulfur puncta inside CD-loaded encapsulins. (**a**) cryo-EM micrographs of CD-loaded (left) and shell-only (right) encapsulin. Electron-dense puncta are only visible in the CD-loaded encapsulin sample localized to the shell interior. (**b**) Representative 2D class averages of CD-loaded encapsulin particles without (top, left) or with (top right) shell subtraction highlighting internal electron-dense puncta. The simultaneous presence of internalized cargo densities (small) and electron-dense puncta (large) can be seen. Representative individual particles of CD-loaded encapsulins containing electron-dense puncta are shown (bottom). (**c**) High-angle annular dark-field imaging (HAADF)-scanning TEM (STEM) micrograph of purified CD-loaded encapsulins. Electron-dense puncta (red arrows) can be seen inside encapsulin shells. (**d**) Elemental mapping via energy-dispersive x-ray spectroscopy (EDS) of representative electron-dense puncta as observed in **c**. HAADF-STEM (left), sulfur map (right). (**e**) Analysis of sulfur valence in empty shell and CD-loaded encapsulin samples determined via x-ray photoelectric spectroscopy (XPS). In the shell-only sample, S(2-) dominates (red) while in the CD-loaded sample S(0) is most prevalent (yellow). (**f**) Cyanolysis of purified empty encapsulin (Enc), free CD, and CD-loaded encapsulin (Enc-CD) samples grown in LB medium to quantify elemental sulfur (S^0^) content. (**g**) High-resolution (HR)-TEM micrograph of a representative electron dense sulfur punctum. Crystal lattice fringes are clearly visible with a d-spacing of 3.4 Å. (**h**) *In vitro* sulfur accumulation assays using Enc-CD at different L-cysteine concentrations. Elemental sulfur content was determined via cyanolysis. (**i**) Elemental sulfur content of purified Enc-CD isolated from cells grown in different defined media conditions. M9 sulfur content: 1.25 mM, M9 -S background: 0.025 mM.

To establish the identity of the observed electron-dense puncta, we set out to determine their elemental composition by performing scanning transmission electron microscopy (STEM) in combination with energy dispersive X-ray spectroscopy (EDS) analysis (Fig. 4c,d and Extended Data Fig. 6b,c). Dark field STEM micrographs showed similar electron-dense puncta inside encapsulin shells as were observed in cryo-EM micrographs (Fig. 4c). Surprisingly, elemental mapping via EDS indicated that puncta contained only sulfur and no other element (Fig. 4d and Extended Data Fig. 6b,c).

This finding suggested that the observed puncta consist of elemental sulfur, which would be in the biologically relatively uncommon S^0^ oxidation state^19^. To confirm the valence of this sulfur species, we carried out X-ray photoelectric spectroscopy (XPS) which showed that in a CD-loaded encapsulin sample, most of the detected sulfur is S^0^ (S 2_p3/2_ binding energy: 163.9 eV), while in a shell-only sample, the majority of sulfur is present in the 2-oxidation state (S 2_p3/2_ binding energy: 168.9 eV) (Fig. 4e)^48^. To further confirm the puncta as elemental sulfur, cyanolysis of purified samples was carried out to detect S^0^ (Fig. 4f)^49,50^. While CD-loaded encapsulins contained substantial amounts of elemental sulfur, free CD and empty encapsulin shells showed much lower levels of S^0^, indicating that elemental sulfur puncta formation is dependent on CD encapsulation inside the encapsulin lumen. Elemental sulfur occurs in different allotropes, such as S_2_, S_6_, S_7_, or S_8_, with S_8_ representing the thermodynamically most stable form under physiological conditions^51^. High resolution TEM imaging revealed that many larger puncta were crystalline, exhibiting clear lattice fringes (Fig. 4g and Extended Data Fig. 6d). The observed interplanar distance (d) values ranged from 2.1 Å to 3.4 Å and did not correspond to any sulfur allotrope deposited in the Crystallography Open Database^52,53^. However, the small apparent d values would not be consistent with any of the larger ring-forming allotropes of elemental sulfur. Considering that CDs function by intermittently forming covalently bound persulfide intermediates, with some CDs having been reported to be able to form short enzyme-bound poly-persulfides in the absence of sulfur acceptors under non-reducing conditions^30,54^, it is possible that continued catalytic activity of encapsulated CD would lead to the formation of long poly-persulfide chains. All but the two terminal sulfur atoms in these chains would be in the S^0^ oxidation state. The buildup of poly-persulfide chains in a restricted space inside a stable protein shell could then lead to the formation of elemental sulfur crystals. Assuming unit cell parameters in the 2-4 Å range with one sulfur atom per unit cell, the maximum sulfur storage capacity per encapsulin shell can be roughly estimated by dividing the available luminal volume (luminal volume – volume occupied by cargo = ca. 2,500,000-3,000,000 Å^3^) by the unit cell volume (ca. 8-64 Å^3^). This back-of-the-envelope calculation yields a storage capacity range of ca. 40,000-375,000 sulfur atoms. Even though this represents only a rough estimate, it seems reasonable to assume that the true storage capacity of CD-loaded encapsulins will be within the same order of magnitude as our estimated range.

To further test the dependence of elemental sulfur accumulation on CD activity and L-cysteine concentration, *in vitro* assays were carried out where purified samples were incubated with L-cysteine, followed by cyanolysis to quantitate S^0^ (Fig. 4h). These experiments showed that elemental sulfur content is titratable and increases over time in CD-loaded encapsulin samples whereas free CD or empty encapsulin shells showed no sulfur accumulation (Extended Data Fig. 6e). The extent of sulfur content increase was found to be L-cysteine- and time-dependent (Fig. 4h). At up to a physiologically relevant L-cysteine concentration of 5 mM, S^0^ content increased linearly for all incubation times tested (1h, 3h, 6h, and 18h). For the 1h and 3h time points, higher L-cysteine concentrations, >5 mM or >10 mM, respectively, resulted in an observed decrease of sulfur content. This is likely due to a combination of substrate inhibition and L-cysteine acting as a reducing agent at higher concentrations and short incubation times leading to the dissolution of stored elemental sulfur. For the longest time point (18h) at the highest L-cysteine concentration (15 mM) an increase in the S^0^ content of ∼550% was observed. Based on the measured amount of accumulated elemental sulfur and the concentration of encapsulins used, the average sulfur content per shell as calculated from bulk assays is ∼150,000 sulfur atoms. This is in good agreement with the estimated storage capacity based on the available internal volume of encapsulins. Taken together, these results indicate that CD-loaded encapsulins are able to actively accumulate S^0^ in an L-cysteine- and time-dependent manner.

As the direct visualization and detection of elemental sulfur via EDS inside cells is hampered by the fact that alcohol- or acetone-based dehydration steps are necessary for preparing appropriate thin sections and that elemental sulfur is soluble in alcohols and acetone, we opted to investigate *in vivo* sulfur puncta formation via an alternative route. Specifically, we set out to modulate the intracellular sulfur pool, which should influence the sulfur content of heterologously expressed and purified CD-loaded encapsulins, through *E. coli* growth assays in M9 minimal medium supplemented with different sulfur sources, namely, L-cysteine or magnesium sulfate. We found that high levels of added L-cysteine and sulfate had a marked effect on the elemental sulfur content of purified CD-loaded encapsulins with an increase in elemental sulfur content of ∼230% in the presence of added L-cysteine and ∼120% in the presence of added sulfate compared to standard M9 minimal medium (Fig. 4i). As Enc-CD directly utilizes L-cysteine, it would be expected that providing additional extracellular sulfur in the form of L-cysteine or its oxidation product cystine – which can be directly imported by a number of dedicated transport systems^55,56^ – would have a more pronounced effect on *in vivo* elemental sulfur accumulation compared to the addition of sulfate which would first need to be converted to L-cysteine via the multistep energy-consuming assimilatory sulfate reduction pathway^57^. Overall, these experiments indicate that the formation of elemental sulfur puncta takes place *in vivo* and that these puncta are stable over extended periods of time inside bacterial cells.

### Encapsulation protects elemental sulfur from cellular reducing agents

Free elemental sulfur is usually unstable inside cells due to the reducing conditions of the cytosol and is easily converted to Na_2_S or H_2_S by biological reducing agents^30^. The fact that elemental sulfur puncta in CD-loaded encapsulins are apparently stable and can be purified argues for a mechanism that protects them from the reducing conditions encountered inside cells. We hypothesize that the protein shell may exclude any protein and small molecule reducing agents encountered in bacterial cells from the encapsulin lumen, thus protecting the internalized elemental sulfur from dissolution.

To directly test this hypothesis, we carried out *in vitro* experiments where purified CD-loaded shells containing sulfur puncta were exposed to different reducing conditions, followed by quantitation of S^0^ content. As the *Acinetobacter* Family 2A encapsulin shell only contains relatively small pores, the largest one being the 5-fold pore with a diameter of 6 Å, any protein-based reducing agent is likely too large to access the shell interior. The main small molecule reducing agent in most cells, including *Escherichia* and *Acinetobacter* species, is glutathione (GSH)^58–61^, which based on its size should similarly not be able to easily transit the 5-fold pore (Fig. 5a). Conversely, smaller synthetic reducing agents like dithiothreitol (DTT) would be predicted to access the shell interior more easily. To test this, purified CD-loaded encapsulins containing sulfur puncta were treated with GSH or DTT and incubated for 3 h, followed by S^0^ content determination (Fig. 5b). Addition of 10 mM GSH, representing the high range of reported cellular GSH concentrations in bacteria^62^, did not decrease S^0^ content significantly, while using 10 mM DTT removed ∼90% of S^0^ from the sample (Fig. 5b). Increasing GSH concentration to a non-physiological level of 20 mM led to a ∼50% decrease in S^0^ content. To further test the protective effect of the encapsulin shell under more native conditions, we incubated purified sulfur-containing encapsulins for 3 h in clarified *A. baumannii* AB0057 lysate (Fig. 5c). Due to the inherent complexity of clarified lysate, background S^0^ signal from lysate was relatively high. We found that the concentration of S^0^ did not substantially change upon lysate incubation and was similar to incubation in buffer containing no reducing agents. In particular, taking the lysate S^0^ background signal into account, S^0^ content in the lysate-incubated sample decreased by less than 15% compared to the buffer-incubated sample. These experiments again suggest that the elemental sulfur puncta observed inside encapsulin shells are stable under reducing conditions as encountered inside cells.

**Fig. 5:**
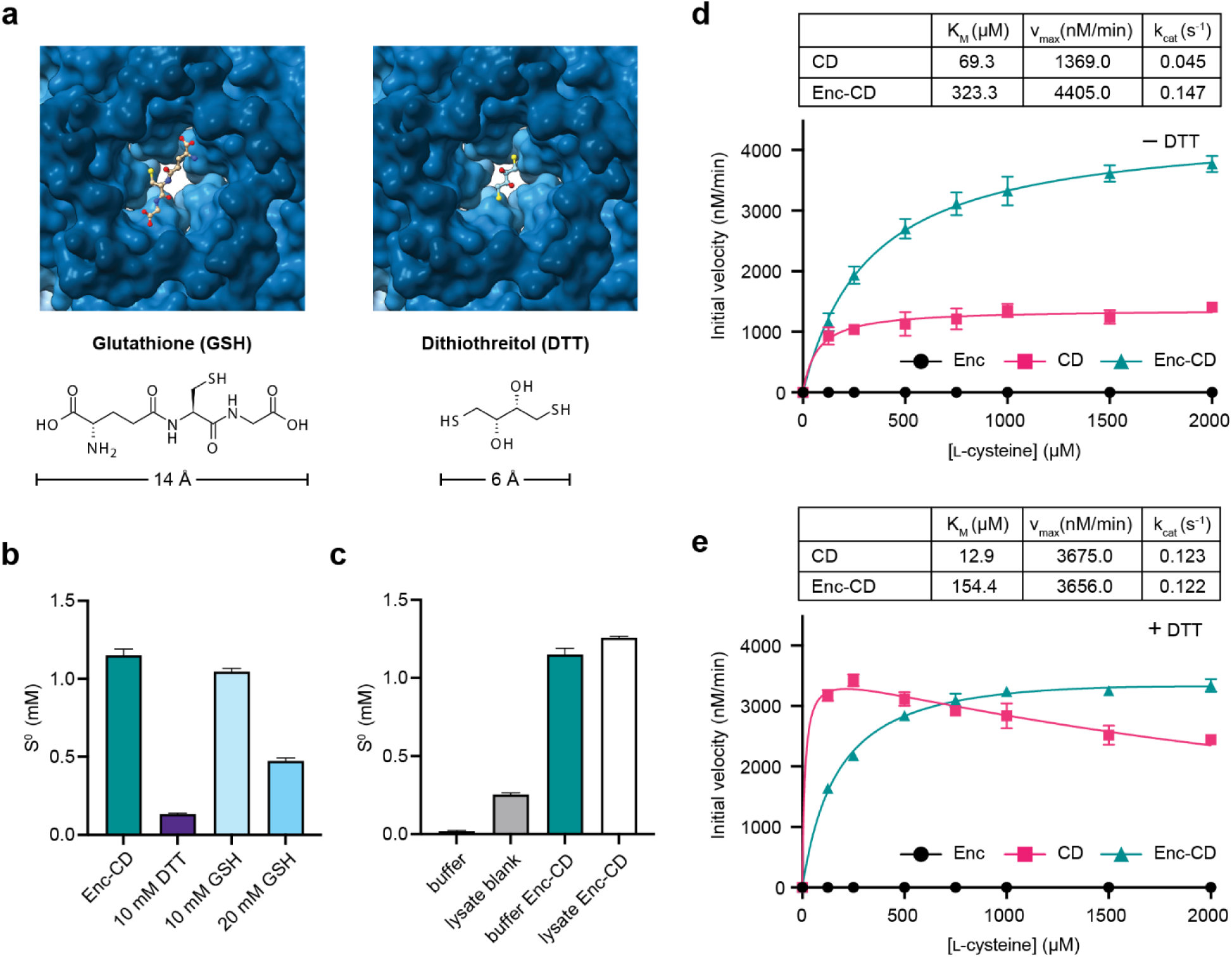
Protection of internalized elemental sulfur puncta from reducing agents and kinetic analysis of desulfurase activity. (**a**) Surface representation of the exterior of the 5-fold pore of the encapsulin shell. At its narrowest point, the 5-fold pore is 6 Å in diameter. For size comparison, glutathione (GSH) (left) and dithiothreitol (DTT) (right) are shown close to the entrance of the 5-fold pore. GSH and DTT are shown in ball-and-stick representation. The chemical structures and approximate longest dimensions of GSH and DTT are shown below. (**b**) Cyanolysis of CD-loaded encapsulins containing elemental sulfur puncta after exposure to reducing agents for 3 h to determine S^0^ content. Enc-CD: Purified CD-loaded encapsulin without exposure to reducing agents. (**c**) Incubation of CD-loaded encapsulins containing elemental sulfur puncta in *Acinetobacter baumannii* AB0057 lysate or non-reducing buffer. (**d**) Saturation kinetics of free CD and encapsulated CD (Enc-CD) in the absence of a thiol-containing sulfur acceptor (DTT, dithiothreitol). Determined kinetic parameters are shown (top). The empty encapsulin shell is used as a negative control. (**e**) Saturation kinetics of free CD and Enc-CD in the presence of DTT as the sulfur acceptor. Kinetic parameters are shown (top). The empty shell is used as a negative control.

### Encapsulation increases CD activity and longevity

We next sought to investigate how encapsulation influences the catalytic activity of CD. For CD kinetic analysis, an established coupled enzyme assay was used where the CD reaction product L-alanine is quantitatively converted to pyruvate by an NAD^+^-dependent alanine dehydrogenase while monitoring NADH production at 340 nm as the reaction readout^23^. Encapsulated CD exhibited increased activity over free CD in the absence of any sulfur acceptor, similar to what was reported for the Family 2A *S. elongatus* system (Fig. 5d)^23^. The k_cat_ for encapsulated CD was determined to be 0.147 s^-1^, more than 3-fold the k_cat_ of free CD (0.046 s^-1^). The observed catalytic activity of our CD was substantially higher than that previously reported for the encapsulated *S. elongatus* CD^23^. When performing CD activity assays in the presence of DTT as a sulfur acceptor, we observed that free and encapsulated CD show similar turnover numbers (free k_cat_: 0.123 s^-1^; encapsulated k_cat_: 0.122 s^-1^) (Fig. 5e). However, free CD exhibited a 5-fold lower K_M_ compared to encapsulated CD and showed marked substrate inhibition. These experiments indicate that CD encapsulation enables high catalytic activity even in the absence of a sulfur acceptor. Even though the mechanistic details are currently unknown, sequestration of up to 12 CD dimers in a constrained nanosized space, like the encapsulin lumen, seems to favor multi-turnover CD activity whereas free CD has to rely on sulfur acceptors to achieve similar kinetics. The relatively high K_M_ value observed for encapsulated CD likely indicates that the shell does, to some extent, act as a barrier for CD substrate binding. This might explain why no substrate inhibition is observed for encapsulated CD where the encapsulin shell would optimize L-cysteine flux to the shell interior.

We further found that catalytic activity after freeze-thawing strongly diminished for free CD, whereas encapsulated CD maintained most of its activity (Extended Data Fig. 7a,b). In the absence of sulfur acceptors, the encapsulated k_cat_ (0.108 s^-1^) only decreased by 25% while no k_cat_ or K_M_ values for free CD could be obtained due to low activity. In the presence of sulfur acceptor, free k_cat_ (0.093 s^-^1) was found to be 3-fold lower than encapsulated k_cat_ (0.026 s^-1^). This suggests that encapsulation of CD may increase enzyme stability and longevity highlighting another potential benefit of CD encapsulation.

## Discussion

In summary, our study demonstrates that a widespread desulfurase-loaded encapsulin nanocompartment represents a novel strategy used by bacteria to store large amounts of elemental sulfur in a crystalline and stable form. Sulfur is essential for all cells and is found in the two proteinogenic amino acids L-cysteine and L-methionine^63^. Sulfur plays a central role in many redox processes and cellular redox homeostasis, for example in the form of protein-bound iron-sulfur clusters or thiol-containing small molecule reducing agents like glutathione^58^. Sulfur is also crucial for the synthesis of important thio-cofactors like coenzyme A, biotin, thiamine pyrophosphate, *S*-adenosylmethionine, and lipoic acid, as well as for many RNA- and protein modifications^35,58,64^. Given the importance of sulfur metabolism, surprisingly many aspects of sulfur and redox homeostasis as well as sulfur trafficking and storage within cells are still unknown.

Our analysis highlights that desulfurase encapsulation inside a protein shell fulfills multiple important roles crucial for the function of desulfurase encapsulins as sulfur storage systems. First, by excluding protein- and small molecule-based cellular reducing agents from the compartment interior, the encapsulin shell creates a distinctive microenvironment that allows the stable storage of elemental sulfur (Fig. 5a-c). This is necessary because elemental sulfur is susceptible to reduction and concomitant solubilization in the reducing milieu of the cytosol^30,63^. Selective shell permeability is achieved through pores optimized for substrate and product transmission while excluding other molecules from the encapsulin lumen (Fig. 2). Second, the pores of the encapsulin shell control substrate flux to the interior of the compartment, essentially controlling the luminal concentration of L-cysteine, thus preventing substrate inhibition observed for unencapsulated CD at higher substrate concentrations (Fig. 5e). Third, encapsulation substantially increases the catalytic activity of CD without relying on an external sulfur acceptor (Fig. 5d). Free CD only reaches comparable catalytic activity when supplied with a thiol-containing acceptor molecule. It appears that CD encapsulation enables the kinetic cycle to close in the absence of a sulfur acceptor. The co-localization and proximity of multiple CD active sites inside the encapsulin shell may allow for inter-CD transfers of persulfides resulting in the build-up of long poly-persulfide chains, eventually leading to sulfur crystal formation (Extended Data Fig. 8). Fourth, the protein shell stabilizes encapsulated CD leading to increased enzymatic longevity (Extended Data Fig. 7). Taken together, the described protein shell-based compartmentalization system represents a unique mechanism for the stable intracellular storage of sulfur.

Our structural analysis of the desulfurase-loaded encapsulin points towards a novel cargo-loading mechanism. CD consists of two domains, a catalytic C-terminal desulfurase domain and a large unannotated N-terminal domain predicted to be intrinsically disordered. Our cryo-EM density reveals multiple likely interaction points of this N-terminal cargo-loading domain with the interior surface of the encapsulin shell. In particular, three discontiguous interactions were identified potentially representing short, conserved motifs within the N-terminal domain. These conserved interaction motifs are connected by long stretches of disordered residues which might be necessary for the correct positioning of the interacting motifs with respect to their binding sites.

The novel encapsulin-based sulfur storage compartment described here stores sulfur in its elemental form, likely as long poly-persulfide chains able to crystallize inside the encapsulin protein shell. We estimate the storage capacity per shell to be on the order of hundreds of thousands of sulfur atoms. No comparable dedicated sulfur storage mechanism has been described before. Select sulfur-oxidizing proteobacterial and Firmicute genera like *Magnetococcus*, *Thiomargarita*, and *Titanospirillum* are able to intermittently form large intra- or extracellular sulfur globules, proposed to be used as a source of energy for cellular respiration and carbon fixation^19,51^. The composition and structure of sulfur granules is still being debated, however, they likely contain elemental sulfur in its thermodynamically most stable form, S_8_, and are surrounded by an irregular protein coat protecting it from immediate dissolution^19^. Most cells, however, do not form sulfur granules and are thought to ‘store’ sulfur in the dynamic L-cysteine pool present in their cytosol. A further potential sulfur storage approach was recently reported in the archaeon *Pyrococcus furiosus* based on uncompartmentalized intracellular thioferrate precipitates, however, the precise function of this system is still unknown^65^. No matter the storage strategy, there needs to be a mechanism to re-mobilize stored sulfur when needed. One component found in ∼50% of desulfurase encapsulin operons that may play a role in sulfur re-mobilization is the rhodanese (Fig. 1c). As a putative sulfur acceptor protein^33–38^, the rhodanese could potentially access stored sulfur through a so far unknown mechanism, followed by the delivery of re-mobilized sulfur to specific downstream processes. In this scenario, the protein shell would represent a specific interaction partner of the co-regulated rhodanese. Such a mechanism could also be coupled with sulfur reduction by a specific thiol-containing small molecule produced under the same conditions as the desulfurase encapsulin and able to enter the encapsulin shell. This may mean that the sulfur stored inside the encapsulin shell represents a privileged sulfur pool that can only be accessed by a specific sulfur acceptor, under specific conditions, and may be destined for specific sulfur-requiring downstream processes. This would be in stark contrast to the easily accessible and promiscuously distributed cellular L-cysteine pool.

The wide distribution of desulfurase encapsulins across many bacterial phyla as well as the diversity of operon structures observed may indicate that different bacteria utilize this sulfur storage system in different ways (Fig. 1). It has been proposed that desulfurase encapsulins may play a role in the canonical sulfur starvation response in freshwater cyanobacteria ^23^. In two further studies, Tn-seq experiments in *Acinetobacter* species showed that the encapsulin shell is either necessary for survival or optimal fitness in different infection models^66,67^.

Taken together, desulfurase encapsulins represent a novel dedicated sulfur storage system in bacteria based on elemental sulfur sequestration inside a defined proteinaceous nanocompartment with potentially different roles across different bacterial species.

## Methods

### Bioinformatic and phylogenetic analysis of desulfurase encapsulins

Family 2A encapsulin sequences were retrieved from UniProt^68^ on June 24, 2022 by searching for non-fragmented encapsulin sequences corresponding to Pfam^69^ PF19307 (Phage capsid-like protein) and excluding Family 2B sequences that were annotated as both Pfam PF19307 and PF00027 (cyclic nucleotide-binding protein), resulting in 1,710 sequences. The Enzyme Function Initiative-Genome Neighborhood Tool (EFI-GNT)^70,71^ was used to generate genome neighborhoods of the identified Family 2A encapsulin-encoding genes. From the resulting collection of Family 2A genome neighborhoods only those encoding genes for Pfam PF00266 (cysteine desulfurase) within 5,000 base pairs of the start codon of the encapsulin-encoding gene were selected. As the Family 2A encapsulin containing Pfam PF19307 also contains phage capsid proteins, phage background needed to be removed. This was achieved by filtering for genes encoding protein sequences corresponding to the Pfam families PF03237 (terminase 6N), PF05133 (phage portal proteins), PF09718 (tape measure proteins), or PF04985 (phage tail proteins). 1,462 Family 2A encapsulin sequences were identified after applying these filters and minor manual curation of the data set (Supplementary Data 1). To generate a sequence alignment for phylogenetic analysis, the encapsulin amino acid sequences were then aligned using MAFFT v7^72^ (MAFFT.cbrc.jp) with default parameters. The sequence alignment was then used to construct a phylogenetic tree by forwarding to the Phylogeny tool on the MAFFT online server with the following parameters: Size = 1,462 sequences x 157 sites, Method = Neighbor-Joining, Model = JTT, Alpha = ∞, Bootstrap resampling = 100. The phylogenetic tree was then assembled into an unrooted radial tree and annotated using iTOL v6^73^.

Disorder prediction of desulfurase cargo proteins was carried out using Disopred3 (Extended Data Fig. 5a)^74^. Sequence logos of conserved motifs found in the N-terminal CLD of desulfurase cargo was created in Geneious Prime 2022.11.

### Molecular cloning, protein production, and protein purification

All expression constructs were ordered as gBlock DNA fragments from IDT (Supplementary Tables S1 and S2). For construction of a C-terminally His_6_-tagged CD, a plasmid containing gBlock RB24 was used as a template to create construct RB30 via PCR using the primers in Supplementary Table S3. For PCR reactions, NEB Q5 DNA polymerase was used. Reactions were prepared according to the manufacturer’s standard protocol, using 10 ng of DNA template. PCR was carried out using a Bio-Rad C1000 thermal cycler. Expression plasmids were constructed via Gibson assembly using linearized pETDuet-1 vector digested with PacI and NdeI and gBlock fragments. Linearized plasmids were gel-purified using an NEB Monarch gel purification kit. Gibson assembly was carried out using Gibson assembly master mix (NEB) according to the manufacturer’s instructions. Plasmids were confirmed via Sanger sequencing (Eurofins). Subsequently, *E. coli* BL21 (DE3) protein production cells were transformed with expression plasmids via electroporation and 25% glycerol bacterial stocks were stored at -80°C.

For encapsulin operon expression and protein production, cells were grown at 37°C, 200 rpm in LB media (10g/L NaCl) with 100 μg/mL ampicillin and induced with 0.1 mM IPTG (encapsulin-containing constructs), or 1 mM IPTG (CD construct) at OD_600_ = 0.6-0.8. Immediately after induction, cells were moved to 18°C for 18-22 h for protein production. Cells were harvested by centrifugation at 5,000 x *g* for 15 min at 4°C, then frozen in liquid nitrogen and stored at -80°C for further use.

Encapsulin protein purification was carried out using a polyethylene glycol (PEG)-based protocol, while CD was purified via Ni-NTA resin affinity purification. All purification steps were carried out at 4°C.

For PEG precipitation-based purification, pelleted cells were resuspended in 20 mM Tris, pH 8 and 150 mM NaCl TBS buffer supplemented with 0.1 mg/mL lysozyme and 0.1 mg/mL DNAse I. Resuspended samples were then lysed by sonication at 75 W, for 5 s/mL, and then clarified by centrifugation at 8,500 x *g* for 15 min. 10% (w/v) PEG 8000 and 600 mM NaCl were added to the supernatant, followed by shaking at 4°C for 30 min. The precipitate was then centrifuged at 8,500 x *g* for 15 min and the resulting pellet resuspended in pH 8 TBS buffer. The resuspended pellet was subjected to gel-filtration chromatography using a HiPrep Sephacryl S500 16/60 HR column with an ÄKTA FPLC system. The resulting encapsulin fractions (identified via light scattering at 320 nm) were buffer exchanged into IEX buffer A (20 mM Tris, pH 8), using an Amicon Centrifugal filter unit with a 100 kDa molecular weight cutoff (Millipore, USA). The resulting sample in IEX buffer A was then subjected to anion exchange chromatography using a HiPrep DEAE 10/16 FF column. Ion exchange was carried by applying a linear salt gradient from 0 to 1 M NaCl. Encapsulin fractions were identified, combined, and subjected to a final gel-filtration step using a Superose 6 10/300 GL column and TBS (pH 8) buffer. All experiments used fresh preparations of protein unless otherwise indicated.

For Ni-NTA affinity purification of the C-terminally His-tagged CD, cells were lysed in 20 mM Tris, pH 8, and 300 mM NaCl buffer supplemented with 0.1 mg/mL lysozyme, 0.1 mg/mL DNAse I, 5 mM β-mercaptoethanol (ME), and SIGMA*FAST* protease inhibitors. Lysis was carried out via sonication at 75 W for 5 s/mL. Samples were then clarified by centrifugation at 8,500 x *g* for 15min. Supernatant was incubated with Ni-NTA resin for 2 h with stirring at 200 rpm and 4°C. Ni-NTA resin was packed into a Bio-Rad Econo glass column, washed with 10 column volumes (CV) of 20 mM Tris, pH 8, 300 mM NaCl, 5 mM ME, and 20 mM imidazole buffer, followed by a 10 CV wash containing 40 mM imidazole in the same buffer. Bound protein was eluted with 10 CV of 20 mM Tris, pH 8.0, 300 mM NaCl, 5 mM ME, and 250 mM imidazole. The His-tag was removed before carrying out any experiment by digestion with TEV protease unless otherwise indicated. TEV protease reactions were carried out in concentrated CD solution, dialyzed against TBS (pH 8.0) supplemented with 5 mM ME overnight. The His-tagged TEV protease was removed from the protein of interest via subsequent batch Ni-NTA purification. Before enzyme assays, purified CD was buffer exchanged into the appropriate buffer to remove residual reducing agent.

### Determination of protein concentration

Protein concentrations were determined using the Pierce Coomassie Bradford Assay kit. Enzyme equivalents of CD in CD-loaded encapsulin (Enc-CD) samples were determined via PLP fluorescence measurements (excitation: 415 nm, emission: 520 nm), using free CD for a standard curve (Extended Data Fig. 7c).

### SDS polyacrylamide gel electrophoresis (SDS-PAGE)

SDS-PAGE was performed using Bio-Rad Mini-Protean 4-20% gels, 1X SDS-PAGE running buffer, 4X Laemmli sample buffer mixed with ME according to the manufacturer’s instructions (Bio-Rad), as well as unstained molecular weight marker (Bio-Rad). Laemmli sample buffer wit ME was brought to 1X concentration by mixing with sample and the resulting mixture boiled at 95°C for 5 min and loaded onto the SDS-PAGE gel. Gels were run at 250 V for 23 min at room temperature. Gels were imaged using the Bio-Rad Chemidoc system in UV stain free mode with auto exposure.

### Negative stain transmission electron microscopy (TEM)

For visualization of encapsulins, samples were applied to a glow discharged Formvar enforced carbon grid (EMS #FCF200-Au-EC) for 1 min, washed with water and dried with blotting paper. Immediately after, 0.5% v/v uranyl formate stain was applied to the grid and immediately blotted away before another 1 min application of 0.5% uranyl formate stain. After final blotting, grids were allowed to dry for at least 15 min before imaging on a Morgagni TEM at 22,000x magnification, with 0.5 s exposure time.

### Native polyacrylamide gel electrophoresis (native PAGE)

Native polyacrylamide gel electrophoresis was performed using Invitrogen NativePAGE 3-12% Bis-Tris gels (ThermoFisher Scientific #BN1003BOX) in 1X NativePAGE running buffer (#BN2007). Samples were prepared using Invitrogen NativePAGE sample buffer (#BN2003). Molecular weight marker (#LC0725) and samples were run at 150 V for 2 h at 4°C, and subsequently stained using Sigma Aldrich Ready Blue protein gel stain (RSB-1L) according to the manufacturer’s instructions.

### Cryo-electron microscopy (cryo-EM) sample preparation, data collection, and data processing

#### Sample preparation

The purified protein samples were concentrated to 3 mg/mL in 150 mM NaCl, 20 mM Tris pH 7.5 buffer. 3.5 µL of protein solution was applied to freshly glow discharged Quantifoil R1.2/1.3 grids and plunged into liquid ethane using an FEI Vitrobot Mark IV (100% humidity, 22°C, blot force 20, blot time 4 seconds, drain time 0 seconds, wait time 0 seconds). The frozen grids were clipped and stored in liquid nitrogen until data collection.

#### Data collection

For Enc-CD, cryo-electron microscopy movies were collected using a ThermoFisher Scientific Titan Krios G4i operating at 300 keV equipped with a Gatan K3 Direct Detector with Bioquantum Imaging Filter. Movies were collected at 105,000x magnification using the Leginon^75^ software package with a pixel size of 0.84 Å/pixel and an exposure time of 3 s, frame time of 50 ms, and total dose of 54 e^-^/A^2^. 5,936 movies were collected for the CD-loaded encapsulin resulting from the heterologous expression of the four-gene operon (Supplementary Table S4). For the encapsulin shell-only sample, cryo-electron microscopy movies were collected using a ThermoFisher Scientific Glacios operating at 200 keV equipped with a Gatan K2 Summit Direct Detector. Movies were collected at 45,000x magnification using the Leginon software package with a pixel size of 0.98 Å/pixel and an exposure time of 5 s, frame time of 200 ms, and total dose of 44 e^-^ /A^2^.

#### Data processing

Data processing for Enc-CD was performed using cryoSPARC v4.2.1^76^. Movies were imported and motion corrected using Patch Motion Correction and CTF fits were refined using Patch CTF. 5,129 movies with CTF fits better than 4.5 Å were selected for downstream processing. Roughly 200 particles were picked manually using Manual Picker and grouped into 10 classes using 2D Classification. Well resolved classes were selected and used as templates for Template Picker to pick particles with a specified particle diameter of 240 Å. 695,842 particles with a box size of 384 pixels were extracted and subjected to 3 rounds of 2D Classification with 100 classes yielding 597,466 particles in good classes. Ab-Initio Reconstruction with 2 classes and I symmetry was carried out next. The main class contained 596,718 particles which were used for further processing. Particles were used as inputs for Homogenous Refinement jobs (with I or C1 symmetry) with the following settings: optimize per-particle defocus, optimize per-group CTF params, and Ewald Sphere correction enabled. The I refinement yielded a 1.78 Å density, whereas the C1 refinement resulted in a 2.18 Å map (Extended Data Fig. 1c,d). Using the Local Resolution Estimation job, local resolution maps for the I and C1 maps were generated yielding similar results (Extended Data Fig. 1e).

### Atomic model building, refinement, and structural analysis

A homologous encapsulin from S. elongatus (PDB ID: 6X8M) was used as an initial starting model for all model building efforts. This starting model was manually placed into the respective cryo-EM map using Chimera v1.14^77^, and was further fit using the Fit to Volume command. The placed monomeric model was then mutated to correspond to the *Acinetobacter baumannii* 118362 amino acid sequence and manually refined against the cryo-EM map using Coot v0.9.6^78^. The resulting model was further refined using Real Space Refine in Phenix v 1.19.2-4158^79^ with default settings and three macro cycles. The model was further refined by iterative rounds of manual refinement using Coot v0.9.6 followed by Real Space Refine in Phenix v1.19.2-4158 until it fit the cryo-EM map satisfactorily. Symmetry restraints were pulled from the map using the Phenix.Find_NCS_from_Map command with I symmetry. The complete shell model was assembled using the Phenix.Build_from_NCS command. This shell model was then used as an input for a final round of Real Space Refine with NCS restraints, three macrocycles, and all other settings set to default (Supplementary Table S4). The model was deposited in the Protein Data Bank (PDB) under PDB ID 8T6R; and the Electron Microscopy Data Bank (EMDB) under EMD-41078.

Channels through encapsulin pores were calculated using the MOLEonline server^80^. The ConSurf Web Server was used to calculate sequence conservation for Family 2A encapsulins and mapping of conservation onto the structure model^81^.

### Scanning transmission electron microscopy (STEM), energy dispersive X-ray spectroscopy (EDS), and high-resolution (HR)-TEM

Enc-CD sample was diluted to 0.1 mg/mL in 20 mM Tris, pH 7.5, and 150 mM NaCl. 4 μL of the protein solution was applied to a glow-discharged Formvar-carbon grid (EMS FCF200-Cu-UB) for 1 min followed by blotting. Imaging was carried out within 24 hours after application to the grid to ensure sample freshness. Both HR-TEM and STEM were carried out using a ThermoFisher Scientific Talos F200X equipped with a Super-X EDS detection system and operated at 200 keV. High-angle annular dark-field (HAADF) images were collected in a range of 56–200 mrad, with a beam convergence angle of 10.5 mrad.

Interplanar distance (d) values were determined using Gatan digital micrograph software. A given crystal with fringes was selected using the ROI tool and Fourier transformed. A mask was applied to opposing spots on the diffraction pattern, and subsequently inverse Fourier transformed. The resulting image was used to produce a profile plot perpendicular to the fringes. The distance between two peaks at similar intensity was measured, and the resulting distance number was divided by N_troughs_ (where N_troughs_ is the number of troughs between peak A and B), yielding the final d-space values.

### X-ray photoelectric spectroscopy (XPS)

XPS measurements were carried out at the Michigan Center for Materials Characterization using a Kratos Ultra DLD system. The sample surface was excited with monochromatic Al kα radiation at 1.486 keV and an incident angle of 54.7°. The source voltage and emission current were tuned to 12 keV and 10 mA, respectively. Broad range survey scans were acquired in the 1200 eV – 0 eV range with a pass energy of 160 eV and step size of 1 eV while the narrow core scans were acquired with a pass energy of 20 eV and step size of 0.1 eV. A charge neutralizer was employed to eliminate any positive charge on the sample surface that is left behind when photoelectrons escape into vacuum. X-rays interacted with the top ∼10 nm in depth of the sample surface and the emitted photoelectrons were directed into an electron energy analyzer to measure their energy. The elemental composition and chemical state of the respective elements was determined using the measured binding energy and intensity of the photoelectron peaks. CASAXPS was used for background subtraction and peak fitting.

### Size determination of sulfur puncta

Size distribution of sulfur puncta was determined from cryo-EM micrographs. ImageJ 1.52p^82^ was used to automatically identify puncta and determine their diameter. All micrographs were first converted to 8-bit before applying a gaussian blur (sigma radius: 2.00). After thresholding, the Analyze Particles function was used yielding a Feret’s diameter distribution representing the size distribution of electron-dense puncta.

### Elemental sulfur content determination by cyanolysis

Elemental sulfur was quantified according to the cold cyanolysis method^49,50^. Briefly, S-S_(n)_ bonds are labile to nucleophilic attack from cyanide (KCN) at basic pH, thus potassium cyanide is mixed with sulfur-containing samples yielding potassium thiocyanate (KSCN), which when mixed with ferric nitrate, produces FeSCN^2+^ which can be detected colorimetrically and quantified at 460 nm.

Samples were diluted to 1 μM CD concentration using TBS buffer (pH 8), unless otherwise indicated. For cyanolysis, a 25 μL sample was mixed with 20 μL of 1 M ammonium hydroxide, 180 μL water, and 25 μL of 0.5 M KCN, in that order. Cyanolysis was carried out at room temperature for 1 h in the dark, after which 5 μL of 37% formaldehyde was added along with 50 μL Goldstein’s reagent. Color was allowed to form for 5 min, then samples were centrifuged at 20,000 x *g* for 5 min before UV-Vis analysis at 460 nm.

### *In vitro* sulfur accumulation assays

Reactions containing 1 μM Enc-CD (based on CD enzyme equivalents) were prepared in 20 mM Tris, pH 8, and 150 mM NaCl. Reactions were started by the addition of the indicated concentration of L-cysteine (0-15 mM) and incubated at room temperature for up to 18 h. Reactions were carried out in triplicate. S^0^ content at different time points was determined by cyanolysis. The addition of 1 M ammonium hydroxide was assumed to terminate the reactions.

### Minimal media protein production with different sulfur sources

Minimal media protein production experiments were performed using an M9 minimal media recipe for protein expression (including casamino acids). The final composition of the M9 base media was 1X M9 salts, 0.4% glucose, 0.5% Casamino acids, 1 mM MgSO_4_, 100 μM CaCl_2_, 36 μM FeCl_3,_ 100 μg/mL ampicillin. The final composition of the sulfur depleted M9 (M9-S) media was 1X M9 salts, 0.4% glucose, 0.1% Casamino acids, 1 mM MgCl_2_, 100 μM CaCl_2_, 36 μM FeCl_3,_ 100 μg/mL ampicillin. The conditions for protein production were the same as described above. For the M9 + 5 mM MgSO_4_ and M9 + 5 mM L-cysteine experiments, cultures were moved to 18°C for 2 h after induction followed by the addition of MgSO_4_ or L-cysteine. Induced cultures were grown for 20 h at 18°C.

### Reducing agent exposure assays

To determine how reducing agents would influence elemental sulfur puncta, reducing agents DTT or GSH were added to a solution containing 20 mM Tris, pH 8, 150 mM NaCl, and 1 μM Enc-CD, and incubated at room temperature for 3 h, followed by cyanolysis for S^0^ content determination.

To produce clarified lysate from *Acinetobacter baumannii* AB0057, cells were grown in 100 mL of LB (0.5 g/L NaCl) at 37°C, 200 rpm, for 18 h. The following day, cells were pelleted by centrifugation and mixed with 1 mL of lysis buffer as used for CD (omitting ME). The resulting mixture was sonicated and centrifuged as described above. The supernatant from the initial centrifugation step was clarified further with multiple 5 min centrifugation steps at 20,000 x *g*, until no further pelleting was observed. The clarified lysate was flash frozen in liquid nitrogen and stored at -80°C until further use. For assays, enzyme was diluted to 1 μM CD equivalents in lysate instead of buffer, incubated for 3 h at room temperature and analyzed by cyanolysis as described above.

### Cysteine desulfurase (CD) activity assays

Cysteine desulfurase activity was assayed similar to previously published methods^23^ by coupling desulfurase activity to alanine dehydrogenase. Briefly, CD abstracts sulfur from L-cysteine to yield L-alanine, which is consumed by alanine dehydrogenase resulting in the producing of an NADH molecule which can be detected using fluorescence spectroscopy. Assay mixtures contained 20 mM Tris, pH 8, 150 mM NaCl, 5 mM NAD^+^, 0.4 U alanine dehydrogenase, 500 nM CD, 10 mM DTT (as sulfur acceptor, when appropriate) and varying L-cysteine concentrations. Reactions were initiated by the addition of substrate. NADH fluorescence was followed using an H1 Synergy plate reader with excitation/emission wavelengths set to 340 nm and 440 nm, respectively. Initial velocities were determined by first calculating the difference in measured relative fluorescence units at which the increase was linear. Conversion to molarity was done using an NADH fluorescence calibration curve. Finally, kinetic parameters were determined in GraphPad Prism by fitting data plotted as initial velocity as a function of L-cysteine concentration with a Michaelis-Menten model (standard), or a substrate inhibition model when appropriate.

## Extended Data Figures

**Extended Data Figure 1:**
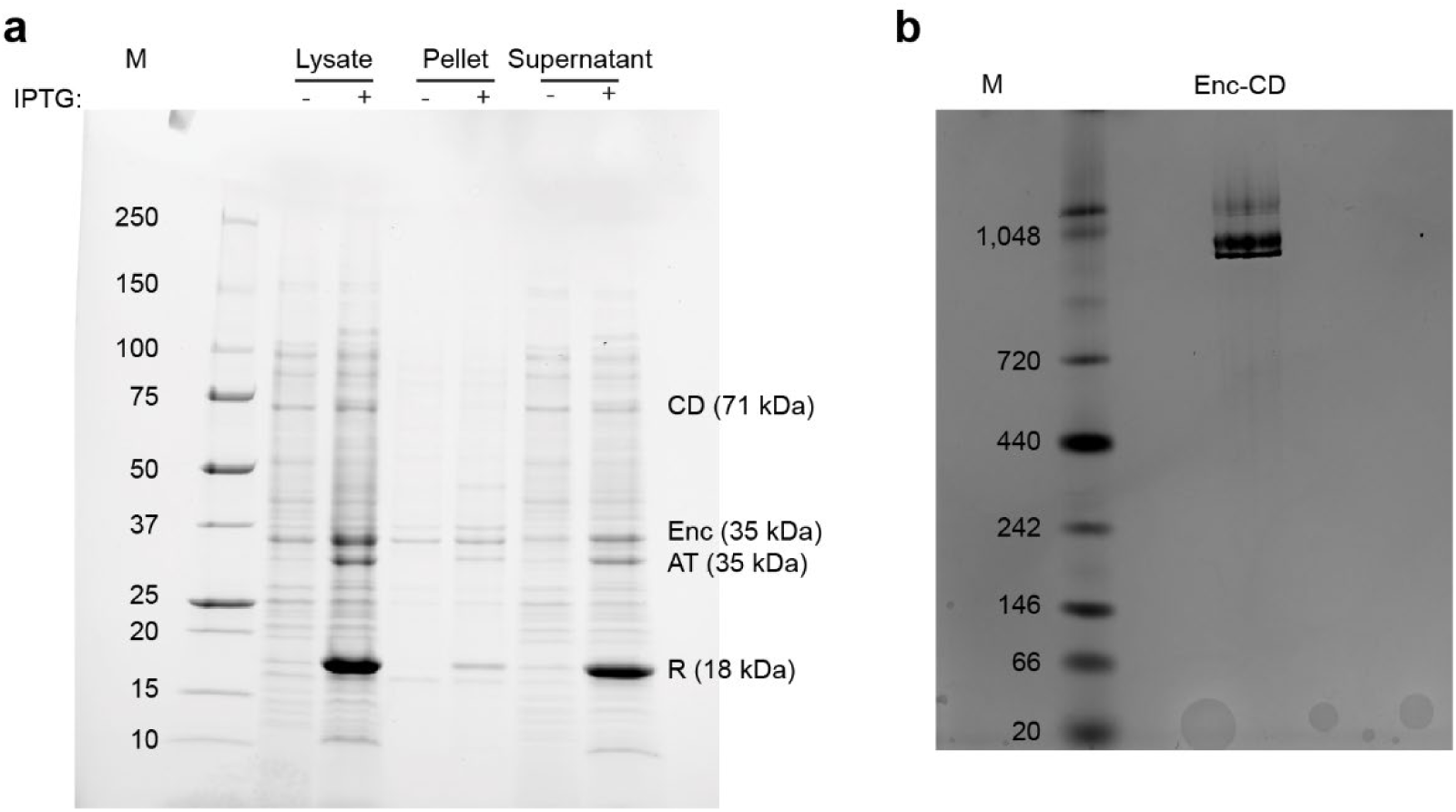
Heterologous expression of the *Acinetobacter* desulfurase encapsulin operon. (**a**) SDS-PAGE analysis of four-gene operon induction test. Rhodanese (R): 18.0 kDa, L-serine *O*-acetyltransferase (AT): 35 kDa, encapsulin shell protein (Enc): 35 kDa, cysteine desulfurase (CD): 71 kDa. (**b**) Native PAGE analysis of purified CD-loaded encapsulin highlighting the lack of a low molecular weight CD band further confirming CD as an encapsulin cargo protein.

**Extended Data Fig. 2:**
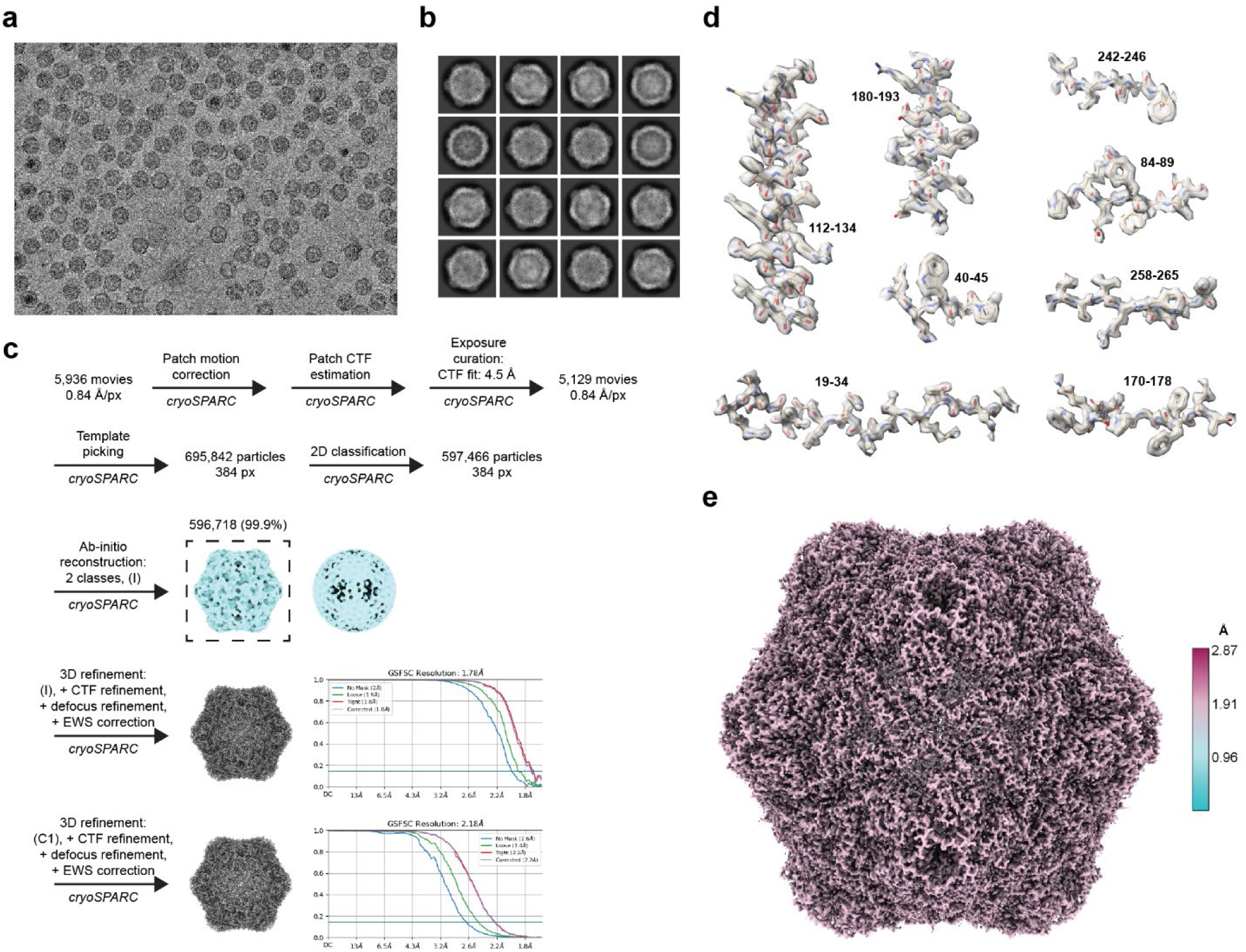
Cryo-EM analysis of the CD-loaded Family 2A encapsulin. (**a**) Representative motion-corrected micrograph of CD-loaded encapsulin resulting from the heterologous expression of the four-gene *Acinetobacter* encapsulin operon in *E. coli* BL21 (DE3). Electron-dense puncta are readily visible. (**b**) Representative 2D class averages of the CD-loaded encapsulin. (**c**) cryo-EM workflow employed in this study. Gold standard Fourier shell correlation (GSFSC) curves for I (1.78 Å) and C1 (2.18 Å) refinements are shown. (**d**) Representative structural elements of the encapsulin shell density after I symmetry refinement highlighting atomic model fit and resolution (1.78 Å). Residue numbering is shown next to each structural element. (**d**) cryo-EM workflow employed in this study. (**e**) Local resolution map of the encapsulin shell after I symmetry refinement. The resolution scale is shown on the right.

**Extended Data Fig. 3:**
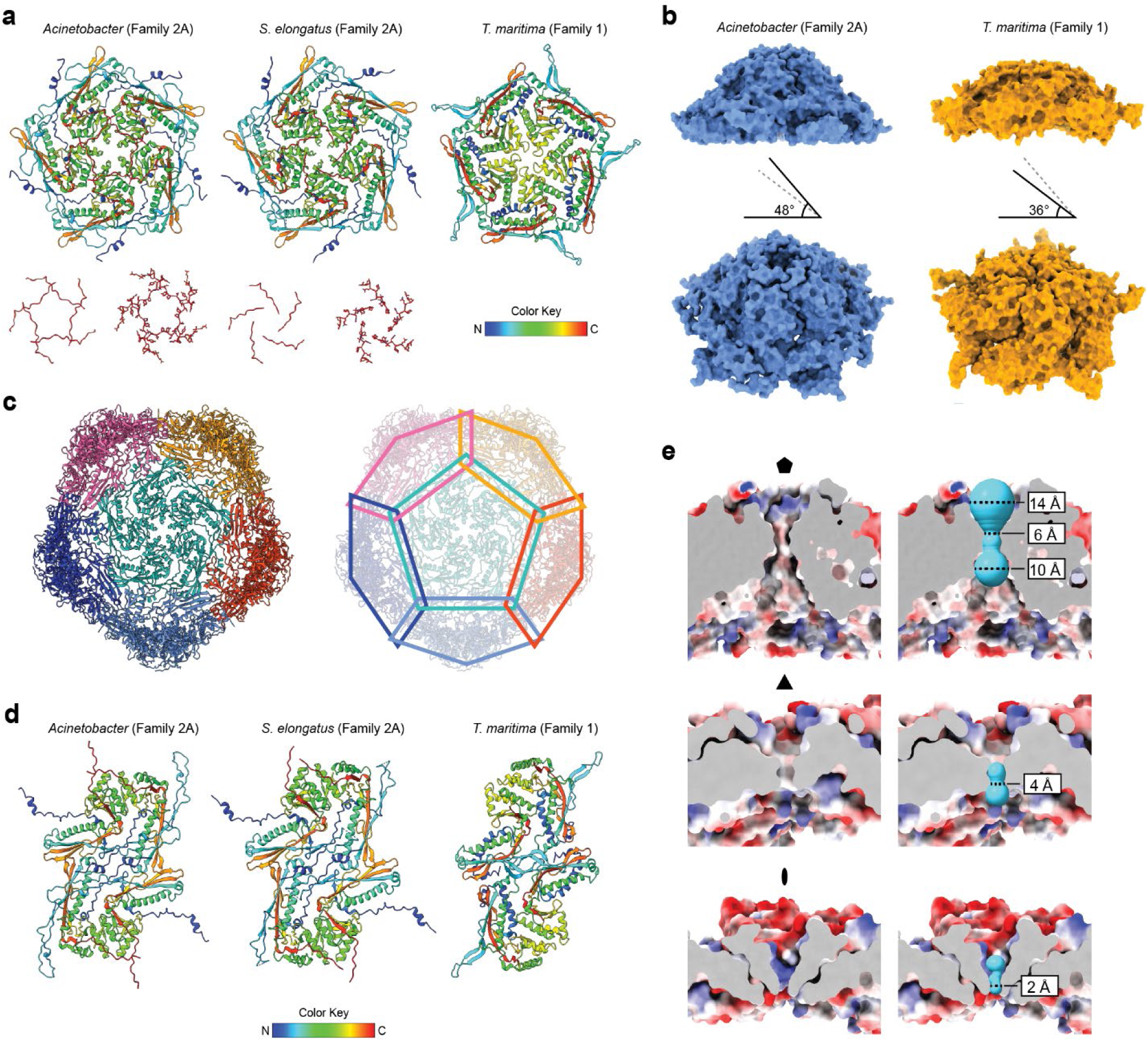
Structural analysis of the encapsulin shell. (**a**) Comparison of the exterior of the 5-fold pores in T=1 Family 2A (*Acinetobacter baumannii* 118362 and *Synechococcus elongatus*) encapsulins and a representative T=1 Family 1 (*Thermotoga maritima*) encapsulin (top). Atomic models shown in ribbon representation using rainbow coloring. The extended C-terminus (red), not present in Family 1, is shown for the Family 2A systems (bottom). In the *Acinetobacter* shell, the C-termini overlap and form an aperture-like structure around the 5-fold pore. (**b**) Side and tilted view of a pentameric vertex for the *Acinetobacter* and representative Family 1 encapsulin. Shell protein angles relative to the 5-fold symmetry axis are shown (Family 2A: 48°, Family 1: 36°) highlighting the 12° extra backward tilt in the *Acinetobacter* system resulting in turret-like vertices. Atomic models are shown in surface representation. (**c**) Chainmail-like topology of the *Acinetobacter* Family 2A shell resulting from the inter-subunit interactions of shell protein N-arm segments. (**d**) Comparison of the exterior of the 2-fold pores in T=1 Family 2A (*Acinetobacter baumannii* 118362 and *Synechococcus elongatus*) encapsulins and a representative T=1 Family 1 (*Thermotoga maritima*) encapsulin (top). Atomic models shown in ribbon representation using rainbow coloring. The differences in subunit interactions at the 2-fold symmetry axis between Family 2A and Family 1 systems are readily apparent. Whereas the E-loops (cyan) of two neighboring subunits directly interact at the 2-fold symmetry axis in Family 1 systems, the N-arms (blue) are responsible for outlining and mostly closing the 2-fold pores in Family 2A systems. (**e**) Side view of the 5-, 3-, and 2-fold pores in the *Acinetobacter* Family 2A encapsulin highlighting pore properties. Channels through pores (right, cyan) were calculated using the MOLEonline server and are shown as surfaces. The narrowest point of each pore is highlighted. For the 5-fold pore, the diameters at the entrance (exterior) and exit (interior) of the pore are shown as well. Atomic models are shown as electrostatic surfaces.

**Extended Data Fig. 4:**
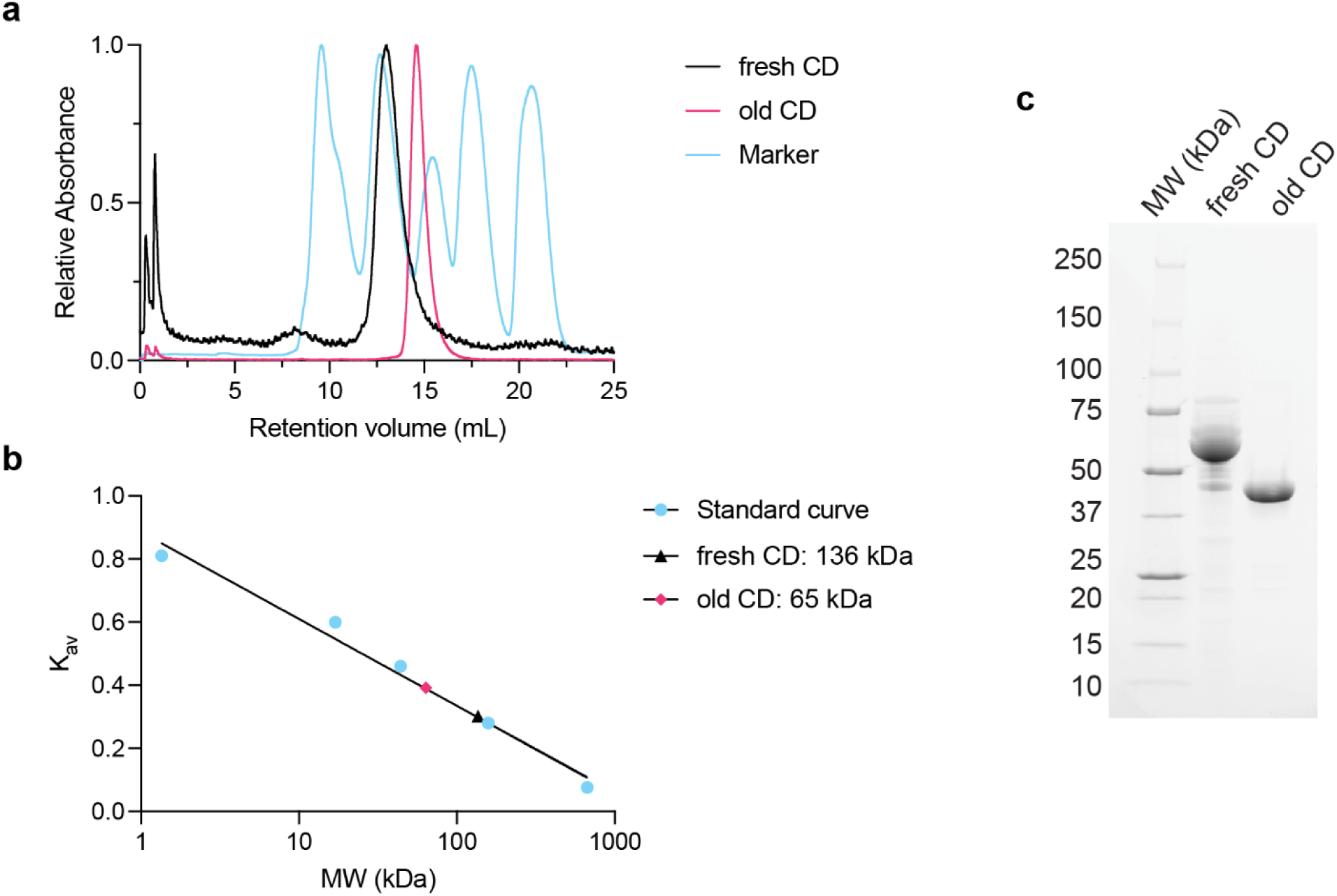
Size-exclusion chromatography of free CD. (**a**) Size exclusion chromatogram of purified free CD using a Superose 6 10/300 GL column. The observed retention time of freshly purified CD (fresh CD) is in line with the expected molecular weight of a CD dimer (142 kDa). CD degrades over time losing the N-terminal CLD yielding a likely dimer with lower molecular weight (old CD). (**b**) Calibration curve used for molecular weight determination based on the following protein standards: thyroglobulin (670 kDa), bovine γ-globulin (158 kDa), chicken ovalbumin (44 kDa), equine myoglobin (17 kDa), Vitamin B12 (1.35 kDa). (**c**) SDS-PAGE gel of fresh and old CD highlighting degradation.

**Extended Data Fig. 5:**
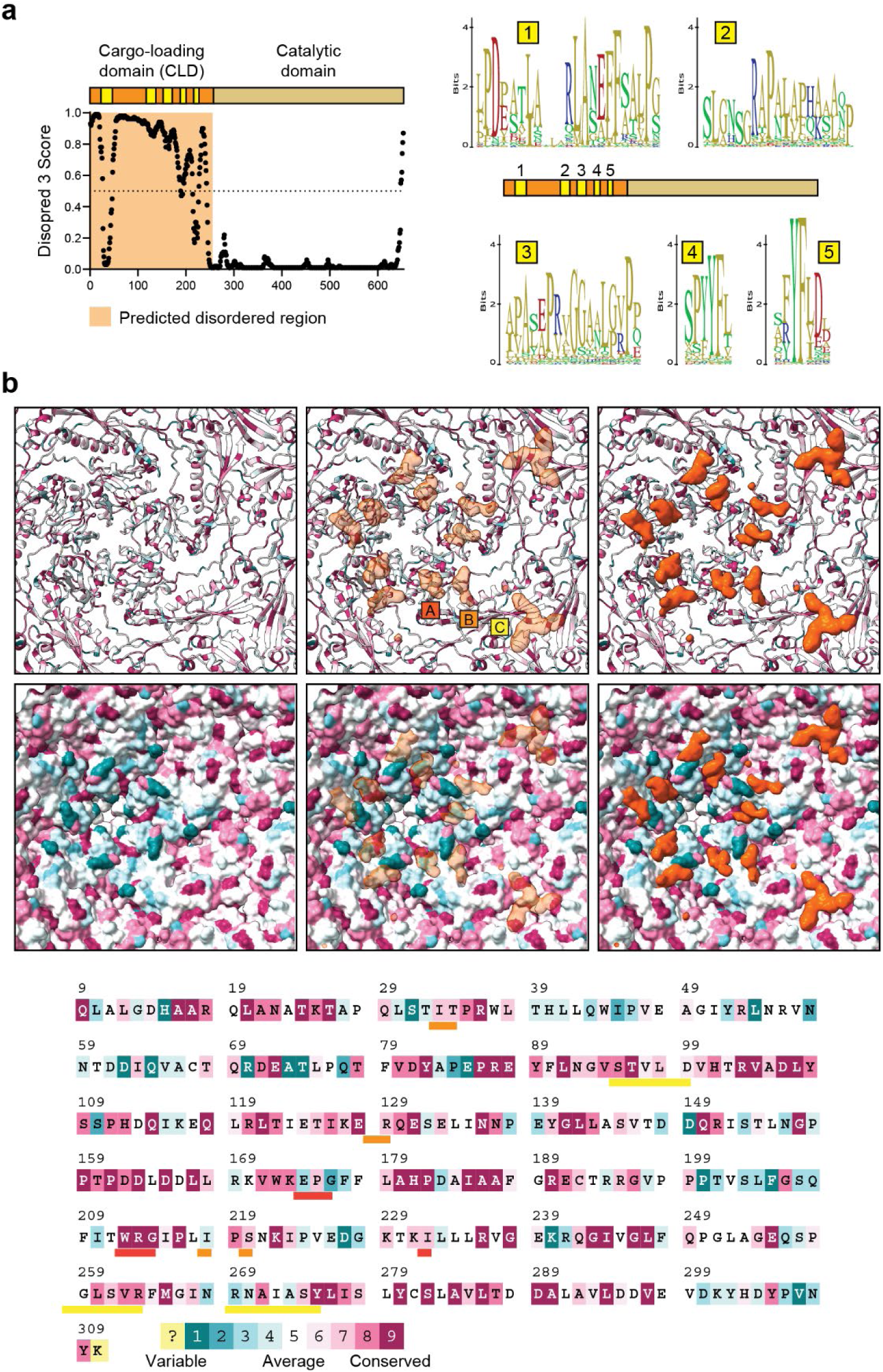
Sequence and structural basis for CD cargo-loading. (**a**) Predicted disorder plot of CD highlighting the disordered N-terminal cargo-loading domain (CLD) calculated using Disopred 3 (top). The first ca. 225 residues are predicted to be mostly disordered with 5 short stretches of sequence predicted to be less disordered corresponding to the 5 conserved sequence motifs identified via CD sequence alignment (below). (**b**) Sequence conservation of Family 2A desulfurase encapsulins mapped onto the atomic model of the *Acinetobacter* shell protein (top). The shell is colored by conservation from variable (teal) to conserved (dark pink) (see below). Both ribbon and surface representations are shown. Additional non-shell cryo-EM densities found to be closely associated with the shell and likely belonging to the CD CLD are shown in orange and localize to the A-domain (around the 5-fold pore) and P-domain (around the 3-fold pore) of each shell protein subunit. The three discontiguous extra densities found for each shell protein subunit are labeled A-C. The *Acinetobacter* shell protein sequence is shown below colored by sequence conservation. Shell residues likely interacting with the three discontiguous CD CLD densities (A-C) are underlined in red (A), orange (B), and yellow (C).

**Extended Data Fig. 6:**
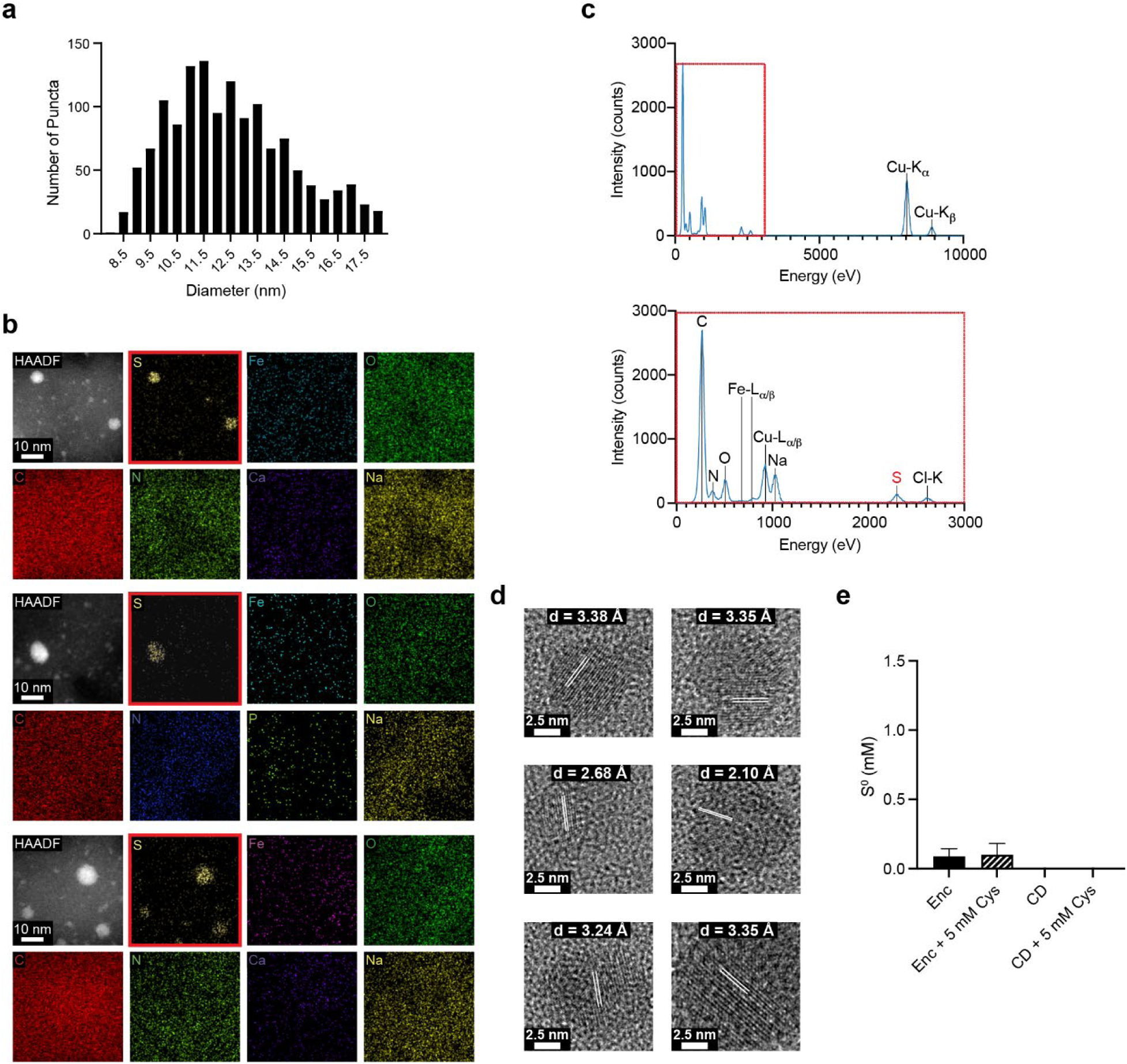
Characterization of elemental sulfur puncta. (**a**) Size distribution of electron-dense sulfur puncta as observed in cryo-EM micrographs. (**b**) Additional representative puncta analyzed via HAADF-STEM and EDS. (**c**) Full EDS spectrum of the area containing two electron-dense puncta shown in Fig. 4d. (**d**) Further representative HR-TEM images of crystalline elemental sulfur puncta. The respective d-spacings are highlighted. (**e**) In vitro sulfur accumulation assay controls. Empty encapsulin shell (Enc) and free CD (CD). Samples were incubated for 3 h followed by cyanolysis to determine elemental sulfur content. No significant sulfur increase was observed.

**Extended Data Fig. 7:**
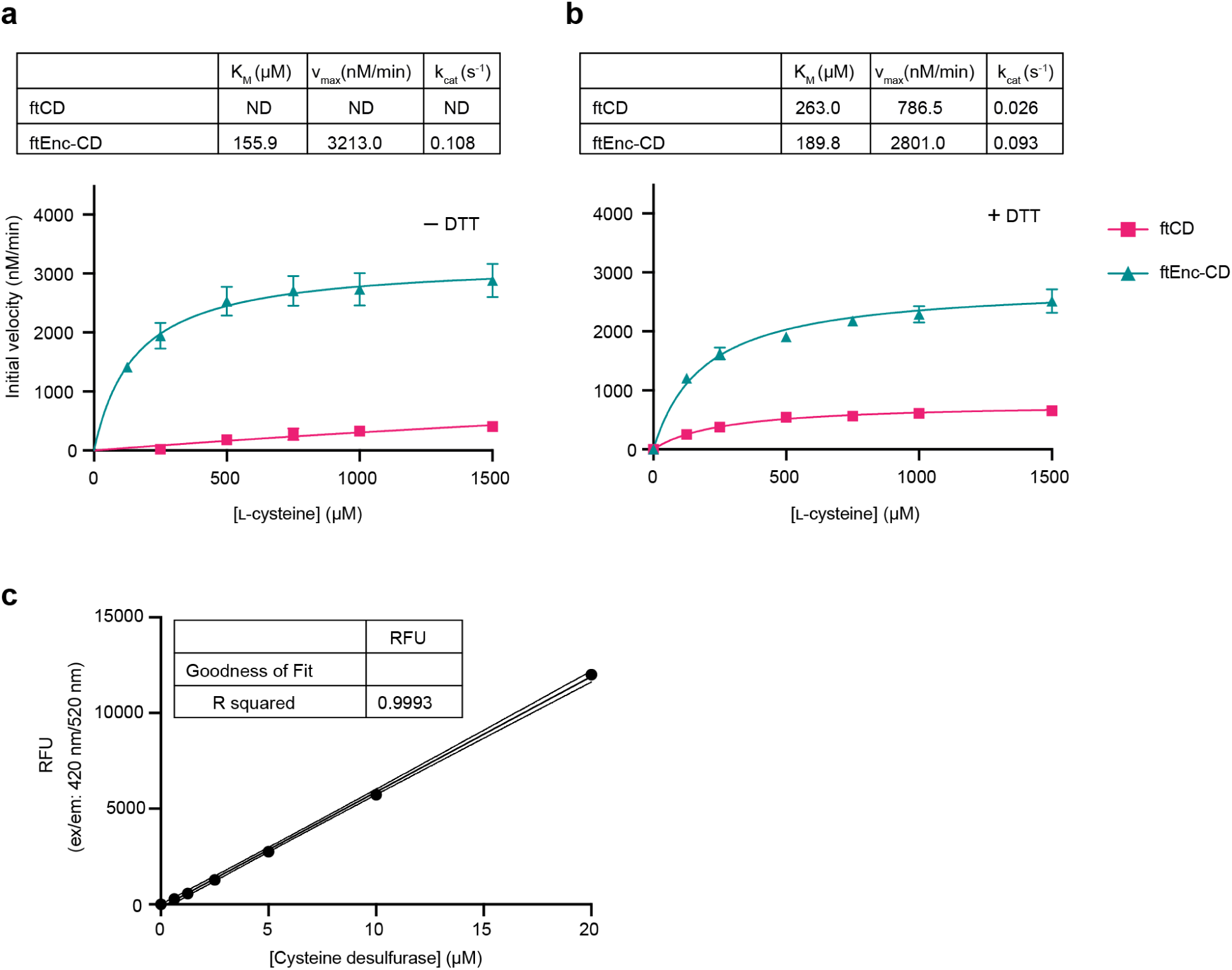
Kinetic analysis of freeze-thawed (ft) samples cysteine desulfurase standard curve. (**a**) Saturation kinetics of freeze-thawed free desulfurase (ftCD) and encapsulated CD (ftEnc-CD) in the absence of a thiol-containing sulfur acceptor (DTT, dithiothreitol). Determined kinetic parameters are shown (top). (**b**) Saturation kinetics of freeze-thawed free desulfurase (ftCD) and encapsulated CD (ftEnc-CD) in the presence of a thiol-containing sulfur acceptor (DTT, dithiothreitol). Determined kinetic parameters are shown (top). (**c**) Standard curve for obtaining enzyme equivalents of encapsulated CD samples based on PLP cofactor fluorescence measurements (excitation: 415 nm, emission: 520 nm) of free CD.

**Extended Data Fig. 8:**
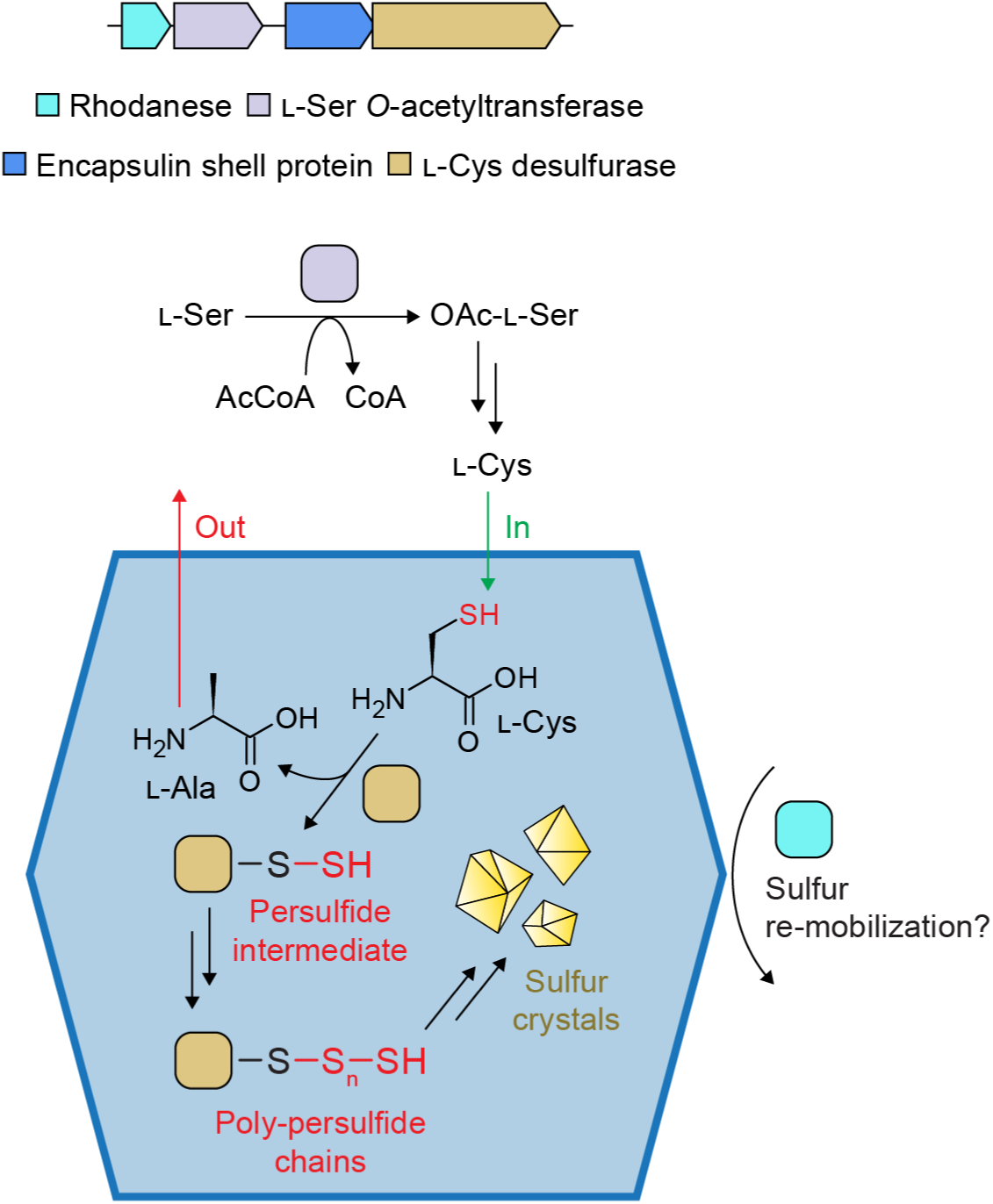
Model of the proposed function of the desulfurase encapsulin-based sulfur storage system. A four-gene operon is shown on top. The proposed functions of the individual operon components are outlined below.

## Acknowledgements

We gratefully acknowledge funding from the NIH (R35GM133325). Research reported in this publication was supported by the University of Michigan Cryo-EM Facility (U-M Cryo-EM). U-M Cryo-EM is grateful for support from the U-M Life Sciences Institute and the U-M Biosciences Initiative. The authors acknowledge the University of Michigan College of Engineering for financial support and the Michigan Center for Materials Characterization for use of the instruments and staff assistance, specifically Tao Ma (HR-TEM, STEM, EDS) and Nancy Senabulya Muyanja (XPS). Molecular graphics and analyses were performed with UCSF ChimeraX, developed by the Resource for Biocomputing, Visualization, and Informatics at the University of California, San Francisco, with support from National Institutes of Health R01GM129325 and the Office of Cyber Infrastructure and Computational Biology, National Institute of Allergy and Infectious Diseases.

## Author contributions

R.B. carried out all molecular biology experiments, protein purifications, in vitro assays and analyses, bacterial growth experiments, and negative stain TEM analyses. M.P.A. performed bioinformatic and phylogenetic analyses, prepared cryo-EM samples and carried out cryo-EM data collection. T.W.G. and M.P.A performed cryo-EM data processing and analysis. M.P.A. carried out atomic model building and refinement. T.W.G, R.B., and M.P.A. wrote the manuscript. T.W.G. directed the project.

