## Supplementary Information for "A widespread proteinaceous sulfur storage compartment in bacteria"

**Supplementary Table S1. Amino acid sequences of proteins produced in this study.**

| Protein | Protein sequence |
|---|---|
| Rhodanese (R) | MNAKLNTETPQSFVSNGSNTSFSAEDILAKAQQYAQEHELNFSGSLSPVDAWQLVQQGEAVLVDV<br>RTNEERKFVGYVPESIHVAWATGTSFNRNPRFLKELESKVGKDKTILLLCRSGNRSTQAAEAAFN<br>AGFEHIYNVLEGFEGDLNEQQQRNQKNGWRIHQLPWQQD* |
| Serine-*O*-acetyltransferase (AT) | MTQWNINAVVQGLQQARHDWRQQQHRTKEFGGRELPSKEALKAILDDLCGILFPMRLGPADLRQE<br>TEDSYIAYTLNRVLTALHAQVQLALNYEAKRHLIHSSPVQQPQELAQTIVQDFANTLPSIRRLLD<br>GDVRAAYEGDPAAHSVDEVLLCYPGIFAIIYHRIAHQLYAQVPLLSRIISELAHSATGIDIHPGA<br>QIGKGFFIDHGTGVVIGETCVIGERVRIYQAVTLGAKRFETNDDGGLKKDYIRHPIVEDDVVIYA<br>GATILGRITIGRGSIIGGNVWLTHSVAAGSQILQSPNESYQKNHIAG* |
| Encapsulin (Enc) | MAKNTDKAQLALGDHAARQLANATKTAPQLSTITPRWLTHLLQWIPVEAGIYRLNRVNNTDDIQV<br>ACTQRDEATLPQTFVDYAPEPREYFLNGVSTVLDVHTRVADLYSSPHDQIKEQLRLTIETIKERQ<br>ESELINNPEYGLLASVTDDQRISTLNGPPTPDDLDDLLRKVWKEPGFFLAHPDAIAAFGRECTRR<br>GVPPPTVSLFGSQFITWRGIPLIPSNKIPVEDGKTKILLLRVGEKRQGIVGLFQPGLAGEQSPGL<br>SVRFMGINRNAIASYLISLYCSLAVLTDDALAVLDDVEVDKYHDYPVNYK* |
| Cysteine desulfurase (CD) | MSTTQLTTSNLQGSPTGPEVTGVNPPANFPDESTIARLANEFFSAVPSSSSSTSSSGLNLDLPVG<br>APPHPPQPGEPFSALSGRAPNIAFQKSLNPEYPEHIPNLPDYPERRIVASAPVVGGANLPRPPFE<br>PHLPQAAHSEPNVPRIEPNLPTGADESPYYFLNDFSAPNAAEPSVKGVEDNYSVAQSFQLPHGDQ<br>LKSLLMEDRFATNTPTQSSASSAFYFLDLPQSGQSYSHYQPSQPIFVASSAQPLDVHAIRRDFPI<br>LQERVNGRPLVWFDNAATTQKPQRVIDRIAYFYQHENSNIHRAAHELAARATDAYEHARNVVARF<br>ISAPSSKEIIFVRGTTEGINLIAKTWGEQNIGEGDEIIVSHLEHHANIVPWYQLTKKTGAKLRVI<br>PVDNTGQIILEELPKLINERTKLLSITQVSNALGTVTPVAEVIKIAHAKGVRVLVDGAQSVSHIP<br>TDVQALDADFFVFSGHKVFGPTGIGAVYAKPELLESMPVWEGGGNMIQDVTFEQIVYQPAPNKFE<br>AGTGNIADAVGLGAALEYVESLGIHNIARYEHDLLEYGQNALSSVSGLRLIGTAHHKASVMSFTL<br>QGYSTEQVGKALNQHGIAVRSGHHCAQPILRRFGVEQTVRPSLAFYNTTEEIDLLVSVLHQLTRN<br>RRTS* |
| Cysteine desulfurase -TEV-His6 | MSTTQLTTSNLQGSPTGPEVTGVNPPANFPDESTIARLANEFFSAVPSSSSSTSSSGLNLDLPVG<br>APPHPPQPGEPFSALSGRAPNIAFQKSLNPEYPEHIPNLPDYPERRIVASAPVVGGANLPRPPFE<br>PHLPQAAHSEPNVPRIEPNLPTGADESPYYFLNDFSAPNAAEPSVKGVEDNYSVAQSFQLPHGDQ<br>LKSLLMEDRFATNTPTQSSASSAFYFLDLPQSGQSYSHYQPSQPIFVASSAQPLDVHAIRRDFPI<br>LQERVNGRPLVWFDNAATTQKPQRVIDRIAYFYQHENSNIHRAAHELAARATDAYEHARNVVARF<br>ISAPSSKEIIFVRGTTEGINLIAKTWGEQNIGEGDEIIVSHLEHHANIVPWYQLTKKTGAKLRVI<br>PVDNTGQIILEELPKLINERTKLLSITQVSNALGTVTPVAEVIKIAHAKGVRVLVDGAQSVSHIP<br>TDVQALDADFFVFSGHKVFGPTGIGAVYAKPELLESMPVWEGGGNMIQDVTFEQIVYQPAPNKFE<br>AGTGNIADAVGLGAALEYVESLGIHNIARYEHDLLEYGQNALSSVSGLRLIGTAHHKASVMSFTL |

| | |
|---|---|
| | QGYSTEQVGKALNQHGIAVRSGHHCAQPILRRFGVEQTVRPSLAFYNTTEEIDLLVSVLHQLTRN<br>RRTSENLYFQGHHHHHHH* |

**Supplementary Table S2. DNA sequences of gBlock fragments used in this study.**

| gBlock | DNA sequence |
|---|---|
| RB14 | AAGTATAAGAAGGAGATATACAATGGCCAAGAACACAGACAAAGCTCAACTTGCGTTGGGC<br>GATCACGCCGCCCGTCAATTGGCGAACGCAACGAAGACCGCACCTCAATTATCGACAATTA<br>CTCCCCGCTGGTTAACGCATTTATTGCAATGGATTCCCGTGGAAGCTGGGATCTATCGCCT<br>GAACCGTGTAAACAACACTGATGATATTCAGGTCGCGTGTACTCAACGTGATGAGGCCACG<br>CTGCCTCAGACTTTTGTGGACTATGCCCCCGAGCCGCGTGAGTATTTCTTAAATGGTGTCA<br>GCACGGTTTTGGATGTGCATACGCGTGTGGCCGACTTGTATAGCTCTCCACATGATCAAAT<br>CAAAGAGCAGTTGCGTTTAACTATTGAGACGATCAAGGAGCGCCAAGAGTCTGAACTTATC<br>AATAACCCTGAATACGGTTTGCTTGCATCAGTGACGGATGACCAACGTATTTCCACTCTTA<br>ATGGCCCTCCAACGCCGGACGACTTAGACGACTTGTTACGCAAAGTGTGGAAGGAGCCTGG<br>GTTCTTCCTGGCGCACCCTGACGCAATTGCTGCTTTCGGCCGCGAATGTACTCGCCGTGGT<br>GTACCTCCCCCAACTGTGTCACTTTTCGGTTCGCAGTTTATTACATGGCGCGGTATCCCAC<br>TGATTCCTTCCAATAAGATTCCCGTAGAAGACGGGAAGACAAAGATTCTGTTATTGCGTGT<br>AGGGGAGAAGCGTCAAGGAATTGTTGGGTTATTTCAGCCGGGCTTGGCGGGGGAGCAGTCT<br>CCAGGGTTAAGTGTTCGCTTTATGGGTATTAATCGTAACGCAATTGCCAGTTATCTGATTT<br>CGTTATACTGTTCATTGGCTGTCCTGACCGATGACGCACTTGCCGTGTTAGATGATGTCGA<br>AGTAGACAAGTACCATGACTATCCCGTGAACTACAAGTAAATTAACCTAGGCTGCTGCCAC<br>C |
| RB16 | AAGTATAAGAAGGAGATATACAATGAATGCGAAGTTAAACACAGAAACGCCCCAAAGTTTC<br>GTATCAAATGGCAGCAACACCTCTTTTAGTGCGGAAGATATTTTAGCTAAAGCCCAGCAGT<br>ATGCACAGGAACACGAATTAAATTTTTCAGGTTCCTTGAGCCCAGTAGACGCGTGGCAACT<br>TGTGCAGCAGGGCGAAGCGGTATTAGTGGATGTCCGTACCAATGAAGAACGTAAATTCGTC<br>GGATACGTCCCGGAATCCATTCACGTCGCCTGGGCCACGGGAACATCATTTAACCGTAATC<br>CACGCTTCTTAAAGGAATTAGAATCGAAGGTAGGAAAAGATAAAACTATCCTTCTTTTATG<br>CCGTAGTGGGAACCGCTCGACTCAAGCTGCGGAAGCTGCTTTTAACGCTGGTTTTGAGCAT<br>ATTTACAACGTTTTGGAAGGTTTTGAAGGGGATTTAAACGAGCAACAACAACGCAATCAAA<br>AGAACGGTTGGCGTATCCATCAGCTTCCATGGCAACAAGACTGATACTTCAATCAATTTTC<br>AGCGAATCAAGGAAAGCTTATGACCCAGTGGAACATTAACGCGGTAGTACAAGGTCTGCAG<br>CAGGCCCGTCACGACTGGCGTCAACAACAGCACCGCACCAAAGAATTCGGGGGGCGTGAGT<br>TGCCGTCGAAGGAGGCGCTTAAAGCTATCTTAGATGACTTGTGTGGGATCTTGTTCCCGAT |

| | |
|---|---|
| | GCGCTTAGGTCCTGCGGACTTGCGTCAGGAAACTGAAGACTCCTATATTGCTTACACTCTG<br>AATCGTGTGTTGACGGCCTTGCATGCACAAGTTCAGCTTGCCCTGAATTACGAGGCAAAAC<br>GCCACCTTATTCACTCCTCCCCAGTCCAACAGCCACAAGAGTTGGCCCAGACTATCGTACA<br>GGATTTTGCGAATACGTTGCCTAGTATCCGCCGCTTATTGGATGGGGATGTCCGTGCCGCA<br>TACGAGGGGGACCCAGCAGCGCACTCCGTCGATGAGGTATTGCTTTGTTACCCCGGTATTT<br>TTGCGATTATTTACCATCGCATCGCTCATCAGTTGTATGCGCAAGTTCCGCTGCTTTCTCG<br>TATCATCTCCGAGTTGGCACACAGCGCCACGGGGATCGATATTCACCCCGGAGCGCAGATT<br>GGTAAGGGATTTTTCATTGACCATGGTACGGGTGTAGTAATCGGAGAAACCTGCGTGATCG<br>GTGAGCGTGTACGCATTTATCAGGCTGTCACTCTGGGAGCCAAACGCTTTGAGACGAACGA<br>TGACGGAGGCCTGAAAAAGGATTACATTCGTCATCCGATCGTAGAGGATGACGTCGTCATT<br>TATGCCGGGGCAACGATTTTAGGACGCATTACCATTGGACGTGGAAGCATTATCGGTGGTA<br>ATGTCTGGCTTACTCATTCTGTTGCAGCGGGTAGCCAAATTTTGCAGTCTCCTAATGAGTC<br>ATACCAGAAAAATCACATTGCGGGCTGAAAACTATAGATGTGAAATAAATAGATAAGGAGA<br>TAAATAAACAAAGAACGTGCAGATTAATAAAATGCTTATTTCTAGTTTAAATATATTTTTA<br>TTTAGAAAATAGATAACTCATTAATAAGCTCAGTATTTAAAAAATTTATAACGCGATTTTA<br>ATTTTAAAACCAGAACTAATTCTAGTTTATATGCATCCTAGAATTTTTATTTTCAAAAAGC<br>CAATGCAATAAGGAAAAATGATTCATGGCAAAG |
| RB17 | AAATGATTCATGGCAAAGAATACTGACAAAGCGCAGCTTGCTTTGGGGGATCATGCCGCCC<br>GTCAACTGGCAAACGCAACAAAAACTGCCCCCCAGTTGTCGACGATCACTCCACGTTGGCT<br>TACACACTTGCTTCAATGGATTCCGGTCGAGGCGGGAATTTATCGTTTGAATCGTGTAAAT<br>AACACAGATGATATCCAAGTCGCATGTACGCAACGCGACGAGGCGACATTGCCACAAACGT<br>TTGTTGATTACGCTCCAGAACCGCGTGAGTACTTCCTGAACGGAGTAAGCACTGTGTTGGA<br>CGTCCATACCCGCGTAGCGGACCTTTACAGCAGCCCGCATGATCAAATCAAGGAACAGCTT<br>CGTCTTACAATCGAGACCATTAAAGAACGTCAGGAATCGGAGTTGATTAATAACCCCGAGT<br>ATGGCCTGCTTGCTTCTGTCACTGACGATCAGCGCATTTCTACCCTTAACGGACCGCCAAC<br>TCCAGACGACCTGGACGACCTTTTGCGTAAGGTTTGGAAAGAGCCTGGGTTTTTCTTAGCC<br>CACCCTGATGCAATCGCCGCCTTTGGGCGTGAATGCACCCGTCGTGGTGTTCCCCCGCCCA<br>CAGTGTCTTTATTCGGGAGTCAGTTCATTACGTGGCGTGGCATTCCTTTGATTCCGAGCAA<br>TAAAATCCCGGTTGAAGATGGAAAGACAAAAATTTTGTTATTGCGTGTCGGGGAGAAGCGT<br>CAGGGAATCGTTGGATTGTTTCAACCTGGGTTAGCAGGCGAACAATCGCCAGGGTTGTCTG<br>TACGCTTCATGGGCATTAACCGTAACGCAATCGCGTCGTATTTAATCTCTCTTTACTGTAG<br>CCTTGCAGTATTAACAGACGATGCTTTGGCAGTGTTAGATGATGTGGAGGTGGACAAATAT<br>CATGACTACCCAGTTAACTACAAGTAATTTGCAAGGGTCACCAACGGGCCCGGAGGTTACT<br>GGAGTAAATCCCCCGGCGAACTTCCCGGACGAGAGCACGATTGCCCGCCTTGCCAACGAAT<br>TTTTCAGCGCCGTCCCAAGTTCTTCCAGCTCAACGAGTAGTTCCGGGTTAAATCTGGACTT<br>ACCAGTAGGGGCGCCTCCTCATCCTCCCCAGCCCGGAGAGCCGTTTTCTGCACTGTCAGGT<br>CGTGCGCCCAACATTGCATTCCAGAAAAGCCTGAACCCTGAATACCCCGAACATATTCCGA<br>ACTTGCCGGACTATCCCGAGCGCCGTATTGTAGCATCTGCGCCCGTCGTAGGAGGGGCTAA |

| | |
|---|---|
| | TCTGCCCCGTCCGCCCTTTGAGCCTCATTTGCCGCAGGCCGCGCATAGCGAACCCAACGTA<br>CCACGCATTGAACCAAACTTACCGACCGGGGCCGACGAAAGTCCTTACTATTTCTTGAATG<br>ACTTCTCAGCACCCAACGCCGCTGAGCCATCCGTAAAGGGCGTGGAGGACAATTACAGTGT<br>CGCACAGTCATTTCAGCTTCCCCACGGAGACCAGCTGAAATCGTTGCTGATGGAAGATCGT<br>TTTGCAACTAATACGCCGACCCAGAGTAGCGCAAGCAGCGCGTTTTACTTCTTAGATTTGC<br>CACAGAGCGGACAATCTTATTCTCACTACCAGCCGTCACAACCGATTTTTGTCGCTTCATC<br>TGCCCAGCCATTAGATGTGCATGCGATCCGCCGCGATTTCCCTATTTTGCAGGAACGTGTG<br>AACGGGCGCCCCTTAGTCTGGTTCGACAACGCCGCCACAACACAGAAGCCACAGCGCGTCA<br>TCGACCGTATCGCATATTTCTATCAGCATGAAAACAGCAACATTCATCGCGCAGCGCACGA<br>GCTGGCCGCCCGTGCTACCGACGCGTATGAACACGCGCGCAACGTAGTGGCCCGCTTTATC<br>AGCGCTCCATCCTCGAAAGAGATCATCTTCGTGCGCGGAACTACTGAAGGTATTAATCTTA<br>TCGCCAAGACGTGGGGCGAGCAGAACATTGGGGAAGGGGACGAGATTATCGTCTCGCACTT<br>GGAACATCACGCTAACATTGTTCCTTGGTACCAGTTGACGAAGAAGACGGGCGCGAAATTA<br>CGCGTGATCCCCGTAGACAATACAGGTCAAATCATCCTGGAAGAGCTGCCGAAATTAATCA<br>ACGAGCGTACGAAATTGTTATCCATCACACAAGTCAGCAACGCCCTGGGAACTGTGACACC<br>AGTCGCGGAAGTGATCAAAATTGCCCACGCGAAGGGAGTTCGCGTCTTAGTGGACGGCGCG<br>CAATCCGTTTCCCATATCCCGACTGATGTTCAGGCCTTGGATGCTGACTTCTTTGTCTTTT<br>CGGGACATAAGGTGTTTGGACCGACAGGAATTGGCGCTGTATACGCAAAGCCTGAACTGCT<br>GGAGTCCATGCCCGTTTGGGAAGGCGGGGGGAATATGATCCAAGATGTCACTTTCGAACAA<br>ATTGTCTACCAGCCGGCCCCCAATAAATTTGAAGCGGGAACCGGAAATATTGCTGACGCGG<br>TCGGGCTGGGGGCAGCCCTGGAGTATGTAGAGTCACTTGGCATTCACAACATTGCACGTTA<br>CGAGCACGACTTGTTGGAATACGGCCAGAATGCTCTGTCCTCAGTGAGTGGGTTACGTCTG<br>ATCGGCACGGCGCACCATAAAGCGTCTGTTATGTCGTTCACCCTTCAAGGTTATTCTACCG<br>AGCAAGTAGGTAAAGCCCTGAATCAACATGGTATCGCTGTACGCTCAGGACACCACTGTGC<br>TCAGCCAATTTTGCGTCGCTTCGGTGTGGAACAAACTGTCCGTCCGTCGCTGGCTTTTTAT<br>AATACCACGGAAGAAATTGACCTGTTGGTTAGTGTGTTACATCAGCTGACACGTAATCGTC<br>GCACGAGTTAAATTAACCTAGGCTGCTGC |
| RB24 | AAGTATAAGAAGGAGATATACAATGCACCACCACCACCATCACGGAGGCGGAGGCAGCGAA<br>AATCTGTATTTCCAATCGACAACCCAGCTTACAACATCAAATTTGCAAGGATCCCCCACTG<br>GACCTGAGGTCACTGGAGTAAACCCGCCCGCGAACTTTCCGGATGAGAGTACTATCGCTCG<br>CTTGGCCAATGAGTTCTTTTCGGCCGTGCCAAGTTCGTCAAGTTCGACTTCAAGCAGCGGT<br>TTGAATTTGGACCTTCCCGTTGGAGCTCCCCCCCATCCGCCTCAACCCGGGGAGCCCTTTT<br>CTGCTCTTTCAGGTCGTGCACCTAACATTGCGTTTCAAAAAAGTTTGAATCCCGAATACCC<br>GGAGCATATTCCAAACCTGCCGGACTATCCAGAGCGCCGCATCGTCGCGAGCGCCCCCGTT<br>GTAGGTGGGGCAAATTTGCCACGTCCGCCGTTTGAACCGCACCTTCCACAAGCTGCGCACA<br>GTGAGCCCAACGTTCCGCGTATCGAACCAAATTTGCCGACGGGCGCCGACGAAAGCCCATA<br>TTATTTCTTAAATGATTTTAGCGCACCGAACGCCGCGGAACCCAGCGTCAAAGGTGTTGAA<br>GACAATTACTCGGTTGCCCAATCTTTCCAGCTGCCCCACGGTGATCAGTTGAAGTCGCTGT |

| | |
|---|---|
| | TAATGGAGGATCGCTTTGCTACTAACACACCGACCCAATCAAGTGCCTCCAGCGCGTTCTA<br>TTTTCTGGATCTGCCACAATCTGGTCAATCCTACAGCCATTACCAACCCTCACAACCGATT<br>TTTGTCGCATCCAGCGCGCAGCCCCTGGATGTTCACGCCATCCGTCGCGACTTCCCCATCT<br>TACAGGAACGTGTCAACGGACGTCCCTTAGTTTGGTTCGACAACGCTGCCACCACCCAAAA<br>GCCACAGCGTGTCATCGACCGCATCGCATATTTCTATCAACATGAGAACTCCAACATCCAT<br>CGTGCGGCACACGAACTTGCAGCACGCGCTACTGACGCCTATGAGCACGCACGCAATGTTG<br>TTGCACGCTTTATCAGCGCCCCAAGTTCAAAAGAGATTATTTTCGTCCGTGGAACCACAGA<br>GGGAATCAATTTGATTGCAAAAACGTGGGGGGAGCAAAACATCGGGGAGGGTGATGAAATT<br>ATTGTATCTCATCTTGAACATCATGCGAACATCGTGCCATGGTACCAGTTGACTAAGAAAA<br>CAGGCGCTAAGCTTCGCGTAATTCCCGTAGATAATACCGGACAAATCATCTTGGAGGAGTT<br>ACCGAAATTAATCAACGAGCGTACAAAGTTATTGTCGATCACTCAGGTGAGCAACGCATTA<br>GGGACCGTTACCCCGGTGGCGGAGGTTATTAAAATTGCGCATGCAAAGGGTGTACGTGTTC<br>TTGTTGATGGTGCCCAGTCAGTGAGTCATATCCCGACCGATGTTCAAGCGCTGGACGCCGA<br>CTTTTTCGTATTTTCTGGTCACAAGGTTTTCGGTCCCACGGGTATCGGCGCTGTCTACGCA<br>AAACCGGAGCTGCTTGAAAGTATGCCAGTCTGGGAAGGCGGAGGAAATATGATTCAAGATG<br>TCACCTTTGAGCAAATTGTCTATCAACCCGCGCCAAATAAGTTCGAAGCAGGAACGGGAAA<br>TATTGCGGACGCCGTGGGATTGGGAGCGGCTTTGGAGTACGTGGAGTCCCTGGGAATTCAT<br>AACATCGCTCGCTACGAACATGATTTACTTGAATACGGCCAGAATGCGTTATCATCAGTCT<br>CCGGATTGCGTTTGATCGGCACTGCACATCACAAAGCTTCGGTTATGTCGTTTACTCTTCA<br>GGGATATTCGACGGAGCAAGTTGGAAAGGCATTGAACCAACATGGGATTGCTGTACGTTCG<br>GGGCACCACTGCGCCCAACCCATTTTACGTCGCTTCGGTGTTGAACAGACGGTCCGTCCGT<br>CTCTGGCATTCTATAACACCACAGAAGAGATTGATTTGTTGGTTAGCGTTCTTCATCAGCT<br>TACGCGCAACCGCCGTACTTCATAAATTAACCTAGGCTGCTGCCACC |
| RB30 | AAGTATAAGAAGGAGATATACAATGTCGACAACCCAGCTTACAACATCAAATTTGCAAGGA<br>TCCCCCACTGGACCTGAGGTCACTGGAGTAAACCCGCCCGCGAACTTTCCGGATGAGAGTA<br>CTATCGCTCGCTTGGCCAATGAGTTCTTTTCGGCCGTGCCAAGTTCGTCAAGTTCGACTTC<br>AAGCAGCGGTTTGAATTTGGACCTTCCCGTTGGAGCTCCCCCCCATCCGCCTCAACCCGGG<br>GAGCCCTTTTCTGCTCTTTCAGGTCGTGCACCTAACATTGCGTTTCAAAAAAGTTTGAATC<br>CCGAATACCCGGAGCATATTCCAAACCTGCCGGACTATCCAGAGCGCCGCATCGTCGCGAG<br>CGCCCCCGTTGTAGGTGGGGCAAATTTGCCACGTCCGCCGTTTGAACCGCACCTTCCACAA<br>GCTGCGCACAGTGAGCCCAACGTTCCGCGTATCGAACCAAATTTGCCGACGGGCGCCGACG<br>AAAGCCCATATTATTTCTTAAATGATTTTAGCGCACCGAACGCCGCGGAACCCAGCGTCAA<br>AGGTGTTGAAGACAATTACTCGGTTGCCCAATCTTTCCAGCTGCCCCACGGTGATCAGTTG<br>AAGTCGCTGTTAATGGAGGATCGCTTTGCTACTAACACACCGACCCAATCAAGTGCCTCCA<br>GCGCGTTCTATTTTCTGGATCTGCCACAATCTGGTCAATCCTACAGCCATTACCAACCCTC<br>ACAACCGATTTTTGTCGCATCCAGCGCGCAGCCCCTGGATGTTCACGCCATCCGTCGCGAC<br>TTCCCCATCTTACAGGAACGTGTCAACGGACGTCCCTTAGTTTGGTTCGACAACGCTGCCA<br>CCACCCAAAAGCCACAGCGTGTCATCGACCGCATCGCATATTTCTATCAACATGAGAACTC |

| | |
|---|---|
| | CAACATCCATCGTGCGGCACACGAACTTGCAGCACGCGCTACTGACGCCTATGAGCACGCA<br>CGCAATGTTGTTGCACGCTTTATCAGCGCCCCAAGTTCAAAAGAGATTATTTTCGTCCGTG<br>GAACCACAGAGGGAATCAATTTGATTGCAAAAACGTGGGGGGAGCAAAACATCGGGGAGGG<br>TGATGAAATTATTGTATCTCATCTTGAACATCATGCGAACATCGTGCCATGGTACCAGTTG<br>ACTAAGAAAACAGGCGCTAAGCTTCGCGTAATTCCCGTAGATAATACCGGACAAATCATCT<br>TGGAGGAGTTACCGAAATTAATCAACGAGCGTACAAAGTTATTGTCGATCACTCAGGTGAG<br>CAACGCATTAGGGACCGTTACCCCGGTGGCGGAGGTTATTAAAATTGCGCATGCAAAGGGT<br>GTACGTGTTCTTGTTGATGGTGCCCAGTCAGTGAGTCATATCCCGACCGATGTTCAAGCGC<br>TGGACGCCGACTTTTTCGTATTTTCTGGTCACAAGGTTTTCGGTCCCACGGGTATCGGCGC<br>TGTCTACGCAAAACCGGAGCTGCTTGAAAGTATGCCAGTCTGGGAAGGCGGAGGAAATATG<br>ATTCAAGATGTCACCTTTGAGCAAATTGTCTATCAACCCGCGCCAAATAAGTTCGAAGCAG<br>GAACGGGAAATATTGCGGACGCCGTGGGATTGGGAGCGGCTTTGGAGTACGTGGAGTCCCT<br>GGGAATTCATAACATCGCTCGCTACGAACATGATTTACTTGAATACGGCCAGAATGCGTTA<br>TCATCAGTCTCCGGATTGCGTTTGATCGGCACTGCACATCACAAAGCTTCGGTTATGTCGT<br>TTACTCTTCAGGGATATTCGACGGAGCAAGTTGGAAAGGCATTGAACCAACATGGGATTGC<br>TGTACGTTCGGGGCACCACTGCGCCCAACCCATTTTACGTCGCTTCGGTGTTGAACAGACG<br>GTCCGTCCGTCTCTGGCATTCTATAACACCACAGAAGAGATTGATTTGTTGGTTAGCGTTC<br>TTCATCAGCTTACGCGCAACCGCCGTACTTCAGAAAACTTGTATTTTCAGGGACACCATCA<br>TCACCATCATCATTAGATTAACCTAGGCTGCTGC |

**Supplementary Table S3. Primers used for the construction of CD-TEV-His6 (RB31).**

| Primer | DNA sequence |
|---|---|
| 5’ pETDuet1 MCS2-CD-TEV-6His fw | AAGTATAAGAAGGAGATATACAATGtcgacaacccagcttacaacatc<br>(lower case: CD 5’ annealing) |
| CD-TEV-6His-3’pETDuet1 MCS2 rv | GCAGCAGCCTAGGTTAATCTAATGATGATGGTGATGATGGTGTCCCTGAAAATACAAG<br>TTTTCtgaagtacggcggttgcgc<br>(lower case: CD 3’ annealing) |

**Supplementary Table S4. Cryo-EM data collection, refinement and validation statistics.**

| | CD-Enc (EMDB-41078) (PDB ID: 8T6R) |
|---|---|
| **Data collection and processing** | |
| Magnification | 105,000x |
| Voltage (kV) | 300 |
| Electron exposure ($e^{-}$/Å$^2$) | 54 |
| Defocus range (μm) | -0.5 to -1.0 |
| Pixel size (Å) | 0.84 |
| Symmetry imposed | I / C1 |
| Initial particle images (no.) | 695,842 |
| Final particle images (no.) | 596,718 |
| Map resolution (Å) | 1.78 (I)<br>2.18 (C1) |
| FSC threshold | 0.143 |
| **Refinement** | |
| Initial model used (PDB ID) | 6X8M |
| Model resolution (Å) | 2.0 |
| FSC threshold | 0.5 |
| Map sharpening *B* factor (Å$^2$) | -57.8 |
| Model composition | |
| Non-hydrogen atoms | 2380 |
| Protein residues | 302 |
| *B* factors (Å$^2$) | |
| Protein | 28.96 |
| R.m.s. deviations | |
| Bond lengths (Å) | 0.006 |
| Bond angles (°) | 1.129 |
| Validation | |
| MolProbity score | 1.27 |
| Clashscore | 2.73 |
| Poor rotamers (%) | 1.91 |
| Ramachandran plot | |
| Favored (%) | 98.00 |
| Allowed (%) | 2.00 |
| Disallowed (%) | 0 |
